# A modular toolset for electrogenetics

**DOI:** 10.1101/2021.09.10.459750

**Authors:** J. M. Lawrence, Y. Yin, P. Bombelli, A. Scarampi, M. Storch, L. T. Wey, A. Climent-Catala, PixCell iGEM team, G. S. Baldwin, D. O’Hare, C. J. Howe, J. Z Zhang, T. E. Ouldridge, R. Ledesma-Amaro

## Abstract

Synthetic biology research and its industrial applications rely on the deterministic spatiotemporal control of gene expression. Recently, electrochemical control of gene expression has been demonstrated in electrogenetic systems (redox-responsive promoters used alongside redox inducers and an electrode), allowing for the direct integration of electronics with complex biological processes for a variety of new applications. However, the use of electrogenetic systems is limited by poor activity, tunability and standardisation. Here, we have developed a variety of genetic and electrochemical tools that facilitate the design and vastly improve the performance of electrogenetic systems. We developed a strong, unidirectional, redox-responsive promoter before deriving a mutant promoter library with a spectrum of strengths. We then constructed genetic circuits with these parts and demonstrated their activation by multiple classes of redox molecules. Finally, we demonstrated electrochemical activation of gene expression in aerobic conditions utilising a novel, modular bioelectrochemical device. This toolset provides researchers with all the elements needed to design and build optimised electrogenetic systems for specific applications.

## Introduction

Advances in synthetic biology have allowed for the development of a variety of genetic circuit-based devices used for sensing applications(*1*), drug and chemical production(*2*) and material synthesis(*3*). However, deterministic regulation of these systems is limited compared to the precise system control provided by electronic circuits. Integrating the control of electronic circuits with the biocatalytic activities of cells has the potential to create reliable and affordable devices for various biomedical and industrial applications(*4*).

One major method used to control biological systems is the use of inducible gene expression systems, which consist of a promoter and its cognate transcription factor that can either activate or repress the expression of genes in response to external stimuli(*5*). These systems have been utilised in molecular biology for the study of biomolecular systems in a variety of organisms(*6*–*9*). Inducible gene expression systems also serve as key tools in synthetic biology, being used as inputs for genetic logic circuits(*10*), in microbial and cell-free biosensors(*11, 12*), and as control systems for CRISPR-Cas9 genome editing(*13*), to name but a few applications.

Whilst chemically inducible and light inducible (optogenetic) gene expression systems are the most popular, due to their strong activities, simplicity and robustness(*14, 15*), other inducible systems have been identified or engineered that respond to heat(*16*), magnetism(*17*), pressure(*18*), osmolarity(*19*), salinity(*20*) and gravity(*21*). Electrogenetics is a nascent field of research in which electrical and electrochemical stimuli are used to control biological processes, including gene expression(*4, 22*). Electrogenetic systems consist of four components: an electrode, a redox inducer, a redox-transcription factor and its cognate promoter (Fig. 1a). The electrode is held at a set potential that is used to oxidise or reduce the redox inducer, which is a cell-permeable electron mediator. The redox inducer in turn oxidises or reduces the redox-transcription factor, which leads to activation or repression of gene expression from its cognate promoter.

**Fig. 1:**
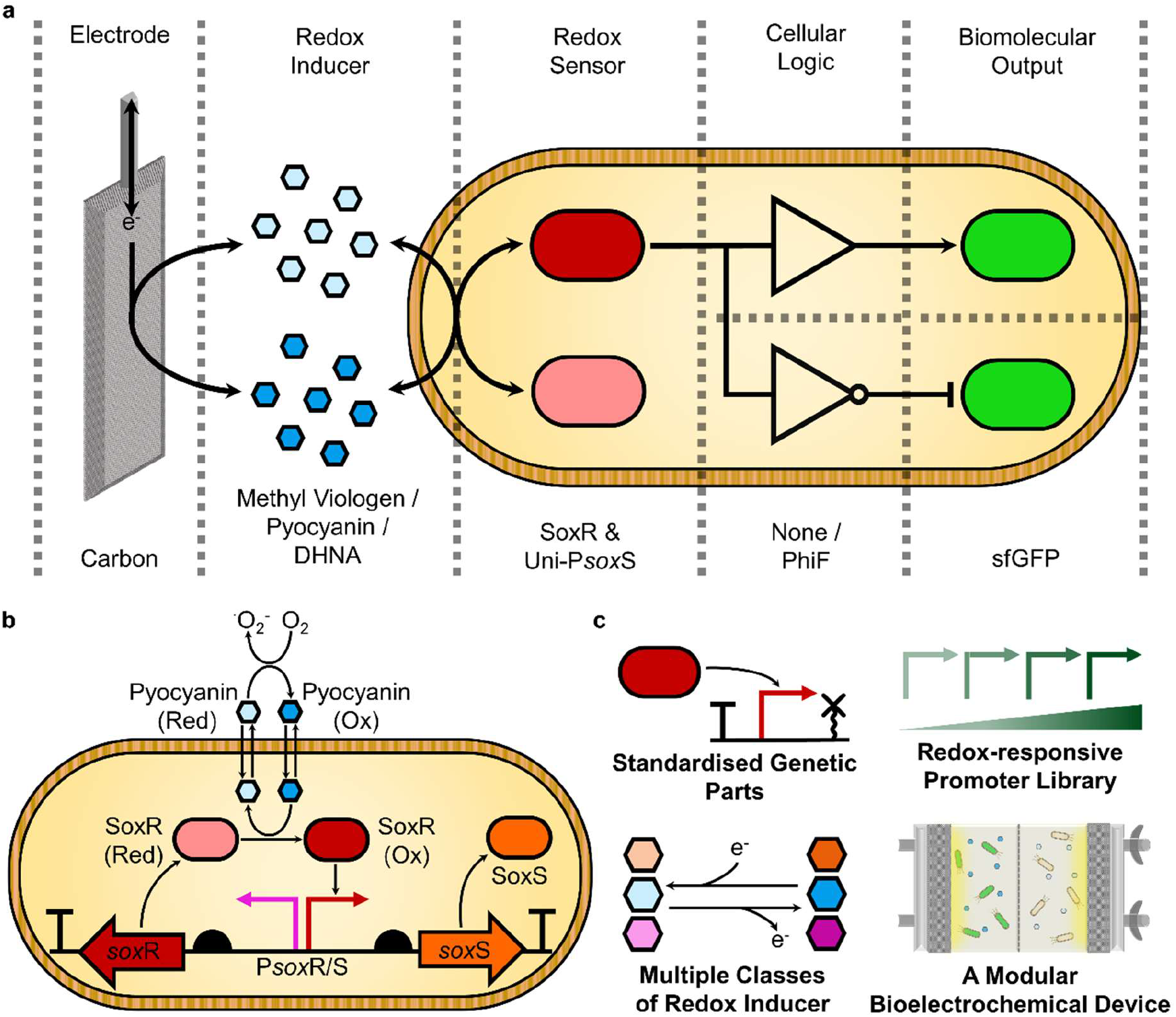
Electrochemical control of gene expression in electrogenetic systems. **a**, Electrogenetic systems consist of a series of electrochemical and genetic modules. An electrode controls the redox state of a redox mediator that the cell responds to with a redox-sensitive transcription factor. Depending on the cellular logic encoded by the genetic architecture used, oxidation of the transcription factor will either activate or repress expression of the chosen gene-of-interest. The identity of specific components tested in this study are labelled beneath the schematic. **b**, The native *sox*RS oxidative stress response system of *E. coli*. SoxR is constitutively expressed from the P*sox*R promoter. Oxidation of SoxR by oxidative-stress inducing redox molecules such as pyocyanin causes transcriptional activation of the P*sox*S promoter. This activation leads to SoxS expression, which activates transcription of numerous genes involved in the oxidative stress response. The redox-dependency of P*sox*S allows for it to be utilised for electrochemical induction of gene expression. **c**, An overview of the genetic and electrochemical tools developed in this study for use in electrogenetics systems.

Unlike other inducible gene expression systems, electrochemical control of gene expression allows for electronic and genetic circuits to be directly connected to one another. The ability to combine electronic controllers easily with cellular biocatalysts is expected to revolutionise bioelectronics, facilitating easy design of devices for biosensing, medical diagnostics and therapies, information processing and energy conversion(*23*–*25*). Electrogenetic systems also have high spatiotemporal control compared to other widely used gene expression systems. They can be used to provide rapid oscillatory gene expression(*22*), as well as spatial gradients of gene expression across cells immobilised in hydrogels(*26*). Already, electrogenetic systems capable of performing digital-to-biological data storage(*27*) and electrically-controlled *in vivo* insulin delivery(*28*) have been developed. Despite these advances, electrogenetics remains an emerging field, with the development of new systems being hindered by a lack of effective and standardised tools, whether they be electrochemical apparatus, redox inducers or genetic parts. Use of the systems has also been limited due to many of them requiring anaerobic conditions(*22*) or complex co-cultures(*29*). Oxidation of redox inducers by oxygen is common, which complicates accurate control of gene expression and can lead to the formation of cytotoxic reactive oxygen species in aerobic conditions.

In a pioneering study, Tschirhart et al. first demonstrated electronic control of gene expression in bacteria by repurposing the *sox*RS oxidative stress response operon of *Escherichia coli*(*22*) (Fig. 1b). In this system a gold electrode was used, with the redox inducer pyocyanin oxidising the SoxR transcription factor to activate the expression of transgenes placed downstream of the P*sox*S promoter. The precise spatiotemporal control of gene expression that has been demonstrated with SoxR-P*sox*S make it a desirable electrochemically inducible gene expression system(*22, 26*). However, the system has not yet been used to activate gene expression controllably in aerobic conditions due to pyocyanin’s reactivity with oxygen(*30*). The native P*sox*S promoter also exhibits a limited dynamic range (*DynR*, the fold-change between the maximum activity and the basal activity) when compared to chemical and optogenetic inducible system(*14, 15, 22*). The lack of a P*sox*S promoter library and the bidirectional nature of the promoter also prevent rational design and standardised assembly of genetic circuits(*31, 32*).

In this work, we redesigned the P*sox*S promoter to facilitate stronger electrochemical induction of gene expression. A unidirectional form of the P*sox*S promoter was engineered with greater dynamic range, allowing for simple construction of genetic circuits. We also created a library of P*sox*S promoters with a spectrum of activities, and tested activation of genetic circuits utilising our promoters with several different classes of redox inducer. Finally, electrochemical activation of gene expression was demonstrated in aerobic conditions using a novel, modular bioelectrochemical device designed for use in electrogenetic systems. These results establish the viability of utilising electrochemically inducible gene expression in synthetic biology. The genetic and electrochemical tools we have developed (Fig. 1c), along with the standardized design principles used, will enable the simple construction of electrogenetic systems for diverse applications.

## Results

### Re-designing the redox-responsive P*sox*S promoter

The bidirectional P*sox*R/S promoter lies at the core of the *sox*RS operon (Fig. 1b; Fig. 2a). The upstream P*sox*R promoter constitutively expresses the SoxR transcription factor, the Fe-S clusters of which can be oxidised by various redox-cycling drugs(*33, 34*). SoxR binds to operator sites in the downstream P*sox*R promoter so that upon its oxidation conformational changes in the protein bring these sites closer together, reducing the distance between the promoter −35 and −10 sites allowing for RNA polymerase recruitment and transcription of the gene for the SoxS regulatory protein(*35*).

**Fig. 2:**
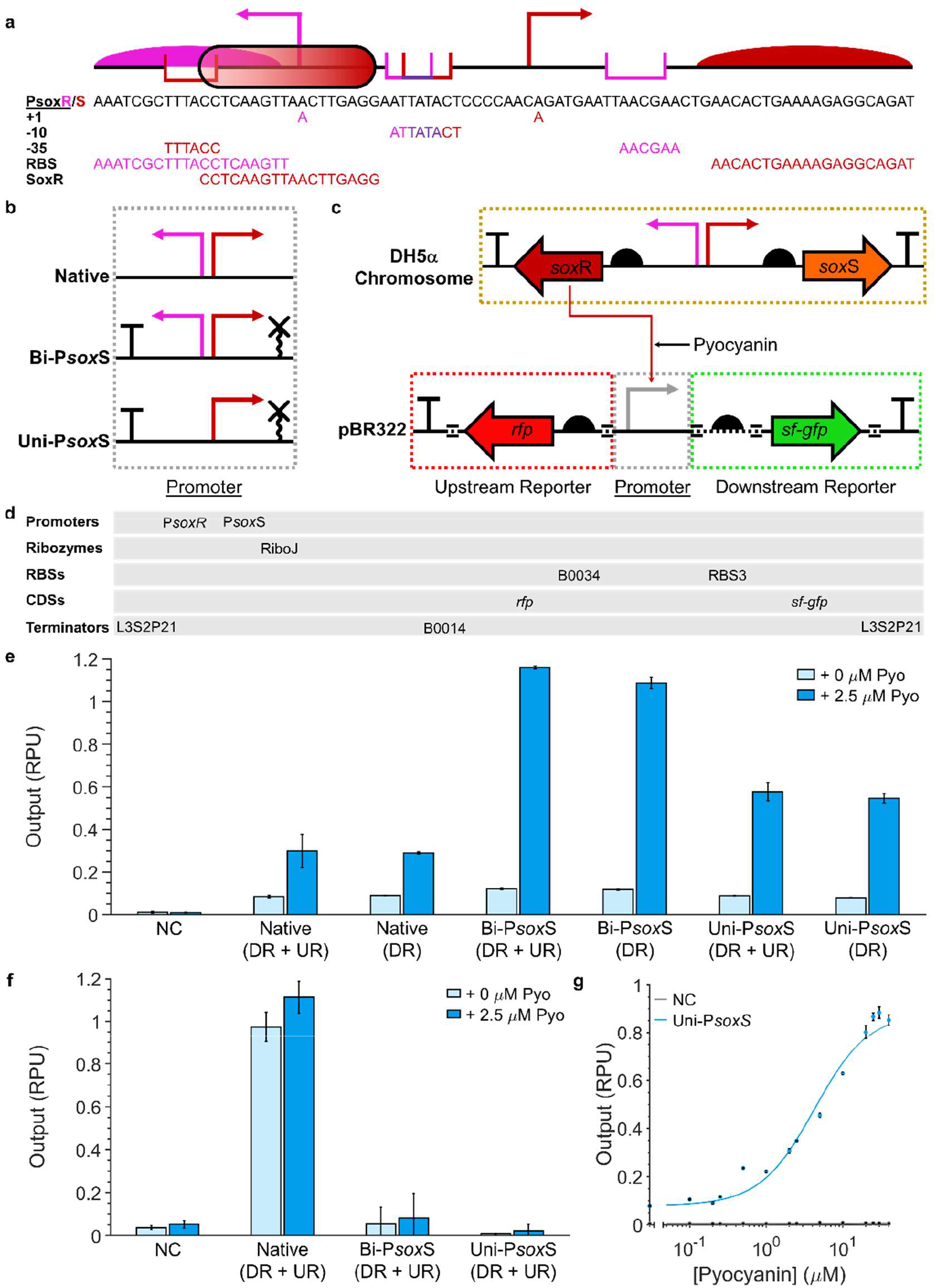
Screening the activity and directionality of mutant P*sox*S promoters. **a**, The sequence of the native P*sox*R/S promoter, with key regulatory components of P*sox*R (pink), P*sox*S (red) and overlapping components (purple) labelled. **b**, Architectures of native and engineered P*sox*S promoters tested. **c**, Genetic circuit design for screening P*sox*S promoters for upstream and downstream transcriptional activity. Circuits can consist of both upstream and downstream reporters, or just the downstream reporter. Black dotted regions of the DNA represent BASIC linkers. **d**, List of genetic parts used. **e**, Downstream activity of engineered P*sox*S promoters with and without 2.5 µM pyocyanin. **f**, Upstream activity of engineered P*sox*S promoters with and without 2.5 µM pyocyanin. **g**, The response function of the Uni-P*sox*S (DR) promoter with pyocyanin. Datapoints represent the mean from three biological replicates with error bars depicting standard deviation (n = 3). UR, upstream reporter; DR, downstream reporter; RPU, relative promoter units.

The bidirectional nature of the P*sox*R/S make it hard to employ in genetic circuits, whilst its overlapping regulatory sequences make it difficult to generate mutant variants of the promoter(*36, 37*) (Fig. 2a). To solve these problems, we have redesigned the P*sox*R/S promoter in order to obtain a unidirectional redox-responsive P*sox*S promoter. We created two variants of the native promoter: Bi-P*sox*S and Uni-P*sox*S. Both were designed with a standardised architecture of an upstream terminator, a spacer sequence and a downstream RiboJ sequence(*38*) (Fig. 2b). These genetic elements were added to reduce promoter context dependency(*39, 40*), whilst the terminator was additionally expected to remove upstream transcriptional activity and aid assembly of the promoter in multi-gene constructs. In Uni-P*sox*S we additionally deleted the P*sox*R −35 regulatory sequence to enhance unidirectional promoter activity. We rationalised that removing P*sox*R activity would also enhance the activity of the promoter by favouring recruitment of RNA polymerase in the P*sox*S direction.

We cloned native, Bi-P*sox*S and Uni-P*sox*S promoter variants into genetic circuits capable of screening their activities through the measurement of fluorescent protein expression with a microplate reader. Downstream and upstream promoter outputs were measured through the stationary phase expression of super-folder green fluorescent protein (sfGFP) and red fluorescent protein (RFP), respectively (Fig. 2c). Fluorescence data was normalised by cell density before, before being divided by the same value recorded from a standardised reference plasmid (Supplementary Table 3) in order to calculate the promoter output in relative promoter units (RPUs)(*41*) (Materials and Methods). This reference plasmid expressed reporters under the control of a J23101 promoter built with the same standardised architecture as Bi-P*sox*S and Uni-P*sox*S. Reporters were expressed under the control of strong synthetic ribosome binding sites (RBSs)(*42*) rather than the native RBS sequences(*37*) in all plasmids (Fig. 2d), and the gene circuits were assembled within a medium-high copy number vector (based on SEVA pBR322)(*43*) to maximise recorded promoter outputs.

*E. coli* DH5α were transformed with either the assembled plasmids containing the promoter variant constructs (Fig. 2b-c) or a negative control plasmid consisting of an empty pBR322 vector (Supplementary Table 3). *E. coli* DH5α contains an intact chromosomal copy of the *sox*RS operon constitutively expressing the SoxR transcription factor (Fig. 2c), allowing for the basal and induced activities of the P*sox*S promoter variants to be measured through the addition of the redox inducer pyocyanin. A pyocyanin concentration of 2.5 µM was selected to maximise promoter outputs without incurring significant cytotoxicity, which we measured by cell growth inhibition (Supplemental Fig. 1).

All promoter variants displayed an induction of downstream promoter output in the presence of pyocyanin. However both mutant variants displayed a significanty enhanced fold-change between the uninduced and induced outputs compared to the native promoter (Fig. 2e). In constructs excluding the upstream reporter, the difference between the induced and uninduced downstream promoter outputs was 3.22-fold for the native promoter, compared to 9.20-fold and 6.86-fold for the Bi-P*sox*S and Uni-P*sox*S variants respectively. Both mutant promoter variants we designed also lacked upstream promoter activity (Fig. 2f), facilitating their use in multi-gene devices without generating unwanted transcripts.

Whilst the fold-change of the Bi-P*sox*S promoter was higher than for the Uni-P*sox*S promoter, the latter has a significant 32.6 % reduction in basal activity in the absence of pyocyanin than the former (Fig. 2e). Consequently, the Uni-P*sox*S variant was selected for use in future constructs. The downstream activity of the promoter was recorded in response to differing concentrations of pyocyanin and was fitted with a response function(*14*) (Fig. 2g; Supplementary Table 1). Response function curve fitting yielded a curve with cooperativity (*n*) of 1.22 and a sensitivity (*K*) of 4.29 µM. This *K* value is comparable to some of the highest performing chemically inducible gene expression systems(*14*).

Previous research has suggested that the P*sox*R promoter is repressed by SoxR(*44*). Furthermore, P*sox*S deletions in the spacer region between the −35 and −10 sites have been shown to convert the promoter from an activator to a repressor in certain strains of *E. coli*(*45*). With this in mind, we designed a unidirectional P*sox*R promoter variant (Uni-P*sox*R), as well as variants of the Uni-P*sox*S promoters with deletions between the −35 and −10 sites with the aim of creating promoters that are repressed rather than activated in the presence of pyocyanin. None of these exhibited significant fluorescence differences when pyocyanin was present (Supplementary Fig. 2) and were therefore not functional repressors in *E. coli* DH5α.

### Development of a redox-responsive promoter library

The deterministic design of complex genetic circuits requires accurate control of gene expression. Because of this, numerous promoter libraries have been generated for *E. coli*, including for both constitutive(*41*) and chemically inducible(*46*) promoters, allowing researchers to tune the expression of genes deterministically within genetic circuits. Here, we have developed a comparable library for Uni-P*sox*S through mutagenesis of the −35 and −10 sites. We selected three −35 sites (termed A-C) and four −10 sites (termed a-d) to test (Fig. 3a) which we expected to provide a range of promoter activities(*46, 47*). The terminal cytosine nucleotides of the −35 site involved in SoxR binding were conserved in all variants(*36*). Uni-P*sox*S variants with different combinations of −35 and −10 sites were screened with the previously described downstream reporter construct (Fig. 2c), using a pyocyanin concentration of 10 µM to induce gene expression. This high concentration was selected to ensure the response of any promoters with low sensitivity would be detected (Fig. 2f). Both the uninduced and induced outputs were measured, with the fold change between them being calculated to estimate the dynamic range. However, it is important to note that some promoters may have lower sensitivities and not be fully induced in the presence of 10 µM pyocyanin, meaning these fold changes may not be an accurate reflection of the dynamic range.

**Fig. 3:**
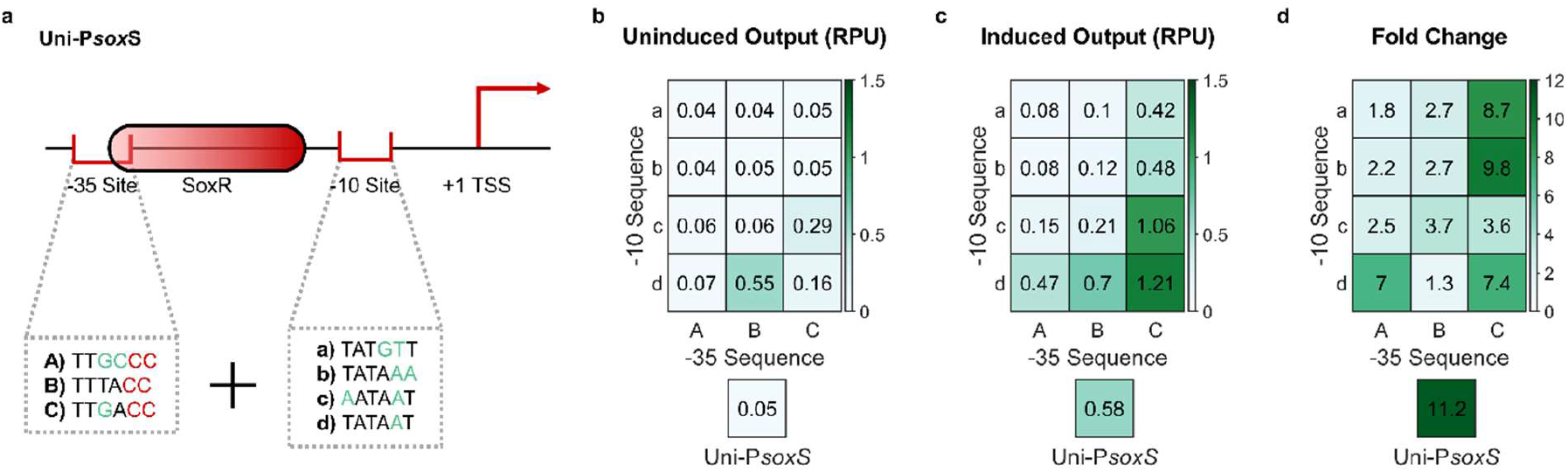
A Uni-P*sox*S promoter library. **a**, Three −35 and four −10 site mutations were screened for tuning Uni-P*sox*S activity. −35 site mutations conserved nucleotides involved in SoxR binding. Promoters were tested using the genetic circuit in Fig. 2c. **b**, OFF activity of the promoter library in the absence of pyocyanin. **c**, ON activity of the promoter library in the presence of 10 µM pyocyanin. **d**, Fold change between ON and OFF activities of the promoter library. Datapoints represent the mean from three biological replicates.

A range of outputs were observed amongst the Uni-P*sox*S mutants (Fig. 3b-d; Supplementary Fig. 3). Whilst no mutations had enhanced fold-change relative to unmutated Uni-P*sox*S, all mutants were inducible and had altered outputs. Some promoters with −10 sites a and b had significant reduction in both the basal and induced output of the promoter, making them preferable for the expression of toxic genes. Promoter mutants Bd, Cc and Cd all had significantly higher basal and induced outputs, with mutant Cd providing the strongest output of any induced promoter at 1.21 RPU, which is comparable to the widely used p*Tet* and p*Lac* chemically inducible promoters(*41*).

### Enhancement and modulation of electrogenetic circuit responses

Whilst the Uni-P*sox*S promoter is functional in *E. coli* DH5α due to its genomic copy of the *sox*RS(*48*), the low level of SoxR expression from the P*sox*R promoter (Supplemental Fig. 2) is not optimised for recombinant protein expression. In order to overcome this low SoxR expression we assembled genetic circuits that not only expressed sfGFP under the control of Uni-P*sox*S in a “output cassette”, but also contained a “sensor cassette” in which SoxR was expressed by a constitutive promoter (Fig. 4a). We developed two versions of these “activator” genetic circuits (Act105 and Act106) with SoxR expression being controlled by a different constitutive promoter in each (Fig. 4b). These constitutive promoters were created using the same standardised promoter design as Uni-P*sox*S (Fig. 2b)(*38*). We tested activator circuits in both DH5α and DJ901, a strain that contains a chromosomal deletion of the *sox*RS operon(*49*). This deletion prevents the strain from expressing antioxidant proteins (*44*) and multi-drug efflux pumps (*50*), which we expected to enhance gene expression induction by pyocyanin, albeit at the expense of increased cytotoxicity(*22*).

**Fig 4:**
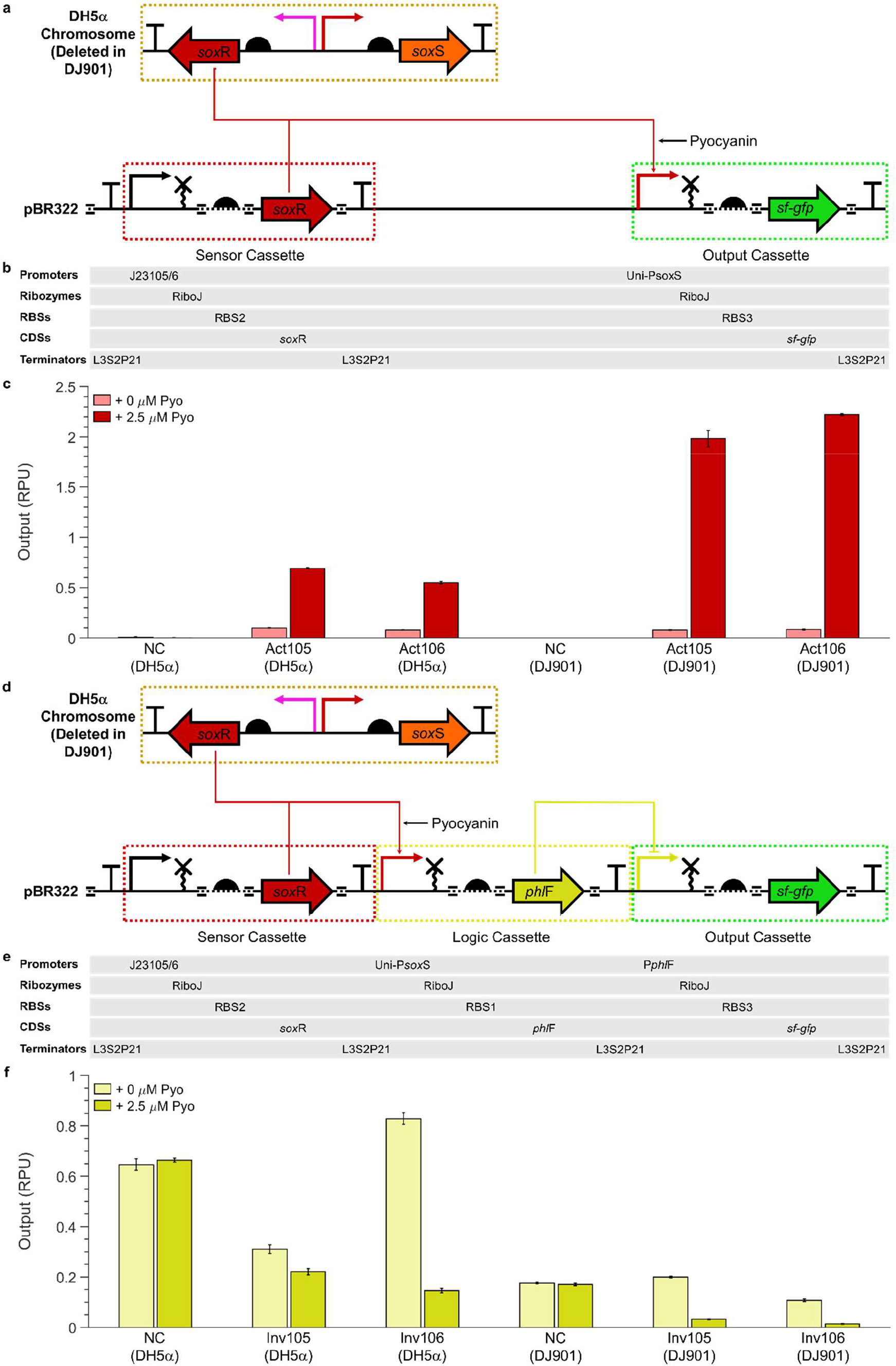
Performance of electrogenetic circuits in different *E. coli* strains. **a**, Genetic circuit design for the activator device. A sensor cassette constitutively expressed SoxR, which, upon oxidation by pyocyanin, activates Uni-P*sox*S gene expression in the output cassette. In DH5α the *sox*RS operon is found on the chromosome, whilst it is missing in the DJ901 deletion mutant. **b**, List of genetic parts used in the activator device. **c**, Activity of activator circuit with and without 2.5 µM pyocyanin. SoxR expression is controlled by J23105 in Act105 and J23106 in Act106. **d**, Genetic circuit design for the inverter circuit. An additional logic cassette is incorporated so that SoxR oxidation by pyocyanin activates expression of PhlF which in turn represses gene expression of the output cassette. DAPG is a chemical inducer of the PhlF-p*Phl*F repression system. **e**, List of genetic parts used in the inverter circuit. **f**, Activity of inverter circuit with and without 2.5 µM pyocyanin. SoxR expression is controlled by J23105 in Inv105 and J23106 in Inv106. Datapoints represent the mean from three biological replicates with error bars depicting standard deviation (n = 3). NC, negative control.

Act105 and Act106 were screened for output in the absence and presence of 2.5 µM pyocyanin. A higher concentration was not used due to the susceptibility of DJ901 to oxidative stress. A negative control construct was also tested (Supplementary Table 3). All circuits demonstrated an enhanced output relative to the negative control, with circuits in DJ901 outperforming those in DH5α (Fig. 4c). Act106 (DJ901) had the best performance, with the highest induced output of 2.22 RPU with 26.3-fold activation being achieved upon addition of pyocyanin, both over twice that observed with Uni-P*sox*S alone in DH5α (Fig. 2e). Interestingly, Act105 and Act106 in DH5α did not provide improved performance relative to Uni-P*sox*S alone in DH5α. Furthermore, whilst the J23106 promoter used in Act106 is much stronger than the J23105 promoter used in Act105 (http://parts.igem.org), the two circuits had comparable outputs. This suggests the activity of activator circuits in DH5α is relatively insensitive to SoxR expression.

Genetic circuit design also allows researchers to modulate the activity of inducible gene expression systems using logic gates. These systems can be simple NOT gates, also known as inverters, which simply invert the signal (change an activator to a repressor and vice versa) or can be more complex logic gates that integrate the signals from multiple input promoters into a single output(*31*). We constructed a second set of genetic circuits, which were similar in design to the activator circuits but included a “logic cassette” in addition to output and sensor cassettes (Fig. 4d). This logic cassette consisted of a PhlF repressor(*14*) being expressed under the control of Uni-p*Sox*S, with sfGFP being expressed from its cognate P*phl*F promoter (Fig. 4e). This logic cassette is a NOT gate, allowing us to repress rather than activate Uni-P*sox*S activity in response to an oxidised redox inducer such as pyocyanin. We therefore termed these “inverter” circuits.

Inverter circuits were tested in the same manner as the activator circuits. The negative control construct consisted solely of the logic and output cassettes, but with the Uni-P*sox*S promoter in the logic cassette replaced with a constitutive promoter. All inverter circuits exhibited a repression of output in the presence of pyocyanin, but to differing extents (Fig. 4f). The outputs of both circuits in DJ901 were very low, with the uninduced state having an output comparable to the basal activity of the activator circuits in the same strain. By comparison, the inverter circuit outputs in DH5α were significantly higher. Whilst this could partially be attributed to the increased strength of Uni-P*sox*S output in DJ901, the reduced output of the negative control in this strain also suggests a differential activity of P*phl*F output between the two strains. As well as the *sox*RS deletion in DJ901, DH5α also contains numerous genomic deletions not found in DJ901 which could contribute to this effect(*48, 49*). Inv106 in DH5α produced the best response, with the highest uninduced output of 0.89 RPU, and a 5.69-fold repression being achieved upon addition of pyocyanin. By successfully demonstrating conversion of the SoxR-P*sox*S system into a repressor we have expanded the applications of electrogenetic systems that utilise it.

### Screening for SoxR Redox Inducers

When designing an electrogenetic system the choice of redox inducer is an important consideration. The membrane permeability of redox inducers will differ between organisms, whilst their midpoint potential and electron transfer chemistry will determine their effectiveness as well as their propensity to partake in cytotoxic or deleterious side reactions such as oxygen reactivity(*30*). Many organisms are also capable of synthesising their own redox inducers, either naturally(*25*) or through genetic engineering(*30*), which can allow for electrogenetic systems to be created without the need for exogenous redox inducers.

The ability of SoxR to be oxidised by different classes of redox inducers provides researchers with flexibility when designing electrogenetic systems(*34*). We demonstrated this flexibility by testing the response of our activator and inverter electrogenetic circuits with five different redox inducers: pyocyanin (a natural phenazine), methyl viologen (a synthetic viologen), DHNA (a natural quinone), riboflavin (a natural flavin) and hydrogen peroxide (Fig. 5g). All of these acted as redox inducers with the Act106 (DJ901) electrogenetic circuit apart from riboflavin and hydrogen peroxide (Supplementary Fig. 5a-c; Supplementary Fig. 4-5). Previous research has demonstrated that *E. coli* lacks riboflavin transporters(*51*), which could explain its lack of induction. The lack of induction by hydrogen peroxide supports previous research that SoxR primarily responds to superoxide-generating, redox-cycling drugs (*52*). The instability of superoxide made it hard to test the response of circuits to superoxide alone, however, previous results have shown that *E. coli* SoxR is relatively insensitive to oxidation by superoxide(*53, 54*).

**Fig 5:**
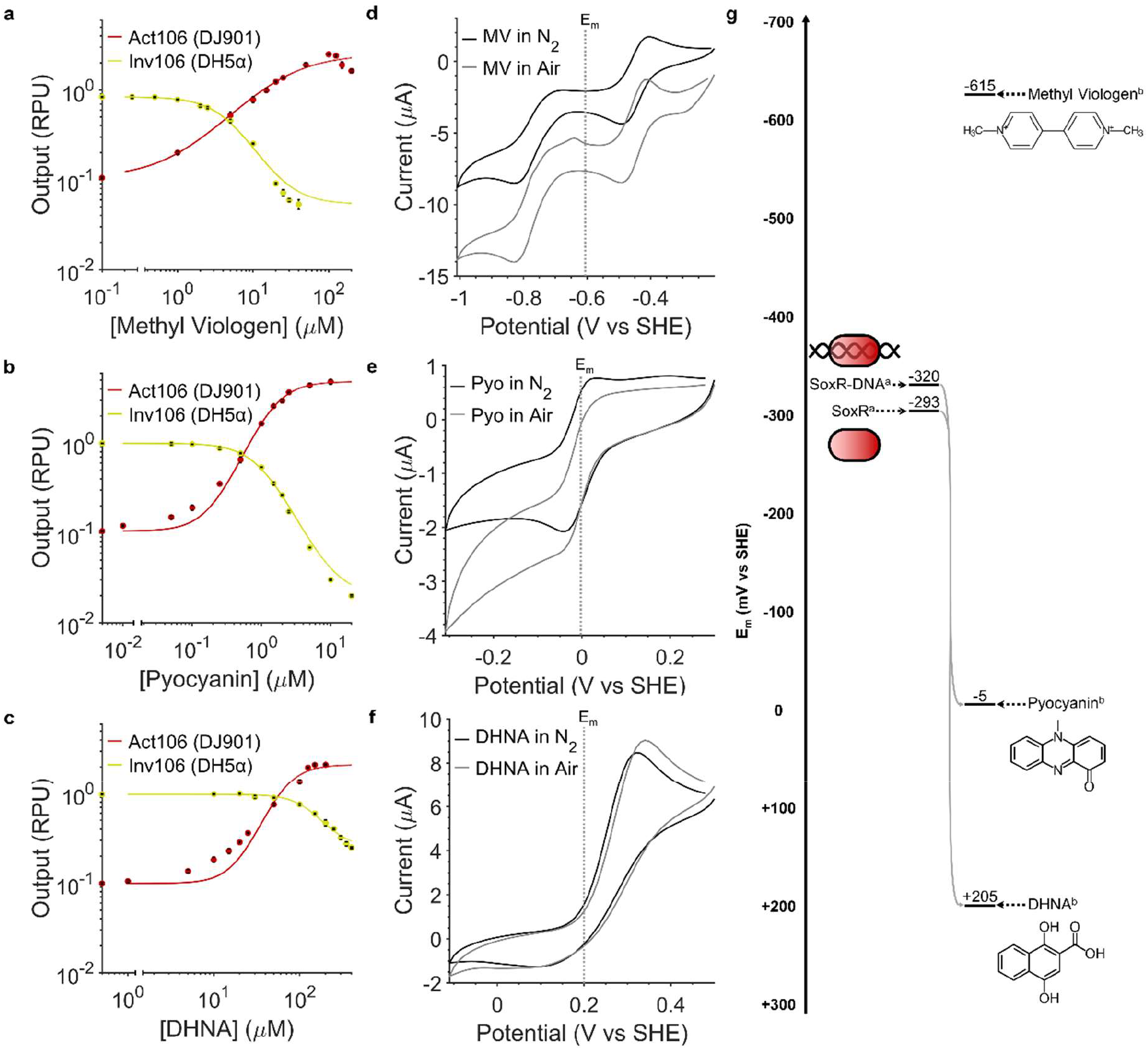
Electrogenetic circuit activation by diverse redox inducers. **a-c** The response function of activator Act106 (DJ901) and inverter Inv105 (DH5α) circuits to methyl viologen, pyocyanin and DHNA. Datapoints represent the mean from three biological replicates with error bars depicting standard deviation (n = 3). **d-f**, Cyclic voltammograms of methyl viologen, pyocyanin and DHNA recorded in aerobic (grey lines) and anaerobic (black line) conditions. LB media was used as electrolyte with 1 mM methyl viologen, 100 µM of pyocyanin or 1 mM of DHNA. The third cyclic voltammetry scan was recorded for each using a scan rate of 10 mV s^−1^. Voltammograms were recorded both in air and in electrolyte purged with N_2_ gas for 30 mins (to remove oxygen) before measurements were recorded with an N_2_ gas stream being maintained in the headspace. **g**, Chemical structures and redox midpoint potentials (E_m_) of redox inducers methyl viologen, pyocyanin and DHNA. Grey arrows indicate the possibility of direct oxidation of SoxR by compounds. E_m_ values from ^a^ Kobayashi, Fujikawa & Kozawa(*33*) and ^b^ this study. Pyo, Pyocyanin.

Response function curves with different redox inducers show pyocyanin provided the highest *DynR* of 46.06 for Act106 (DJ901) and 49.97 for Inv106 (DH5-α) (Supplementary Table 1). These values are comparable with the best performing chemically(*14*) and optogenetic(*15*) inducible gene expression systems available. The maximum output for Act106 (DJ901) was also 4.82 RPU, which is larger than any of the high-performance, plasmid-based, “Marionette” chemically inducible gene expression systems that were produced by directed evolution(*14*). By comparison methyl viologen and DHNA provide a lower, but still workable *DynR* to pyocyanin (Fig. 5a-c; Supplementary Table 1). This improved ability of phenazines to activate the SoxR-P*sox*S system in comparison to viologens and quinones has been reported previously(*54*).

Cytotoxicity of the redox inducers varied (Supplementary Fig. 6). Methyl viologen was the most cytotoxic inducer, providing significant growth inhibition to both Act106 (DJ901) and Inv105 (DH5α) from concentrations of 1 µM. This toxicity is attributable to both its superoxide generating ability and its synthetic nature preventing it from being degraded by cells(*55*). As expected, pyocyanin was much more cytotoxic to Act106 (DJ901) due to the deletion of the entire *sox*RS operon in the strain(*49*), with cell growth being almost completely inhibited at 10 µM with Inv106 (DH5α) exhibiting a <20 % growth inhibition at the same concentration. DHNA interestingly was only significantly cytotoxic to Inv106 (DH5α), which can likely be attributed to differences in the resistance of their respective background strains.

We also tested the performance and cytotoxicity of the Inv106 (DH5-α) in M9 minimal media using pyocyanin as a redox inducer, as previous results have suggested minimal media may provide improved fold-changes with the SoxR-P*sox*S system(*22*) (Supplementary Fig. 7a; Supplementary Table 1). Whilst the value of *K* is much smaller in this condition (0.09 µM in M9 vs 0.57 µM in LB), this improved sensitivity is achieved at the expense of a reduced *DynR* (49.97 in LB vs 38.15 in M9). Little difference in the cytotoxicity of pyocyanin in the two media were observed (Supplementary Fig. 6e & 7b).

Cyclic voltammetry was performed on the successful redox inducers pyocyanin, methyl viologen and DHNA to determine better their electrochemical properties and the mechanism by which they oxidise SoxR (Fig. 5d-f). Midpoint potentials (E_m_), the potential at which a 50 % of the inducer is expected to be oxidised and 50 % reduced, were determined from these voltammograms (Fig. 5g). If an E_m_ value of an inducer is more positive than the E_m_ of SoxR at −320 mV vs SHE (mV henceforth) when bound to DNA (*33*), then that redox inducer is theoretically capable of directly oxidising the transcription factor. The −5 mV E_m_ of pyocyanin and +205 mV E_m_ of DHNA suggest they are both capable of directly oxidising SoxR and activating P*sox*S expression in the process. The −615 mV E_m_ of methyl viologen demonstrates that it is incapable of directly oxidising SoxR and must therefore activate P*sox*S expression by an indirect mechanism, as has been posited by previous studies(*54*).

Voltammograms recorded in air (aerobic) and under N_2_ purging (anaerobic) conditions were also compared to determine the extent of oxygen reduction by these redox inducers. The relative insensitivity of SoxR to superoxide(*33, 54*) and hydrogen peroxide(*52*) means that oxygen reduction by redox inducers is not expected to activate expression of genes downstream of P*sox*S but will instead limit the ability for the redox inducer to be fully reduced by an electrode. Whilst cyclic voltammograms of DHNA showed no shift in current between scans recorded in air and under N_2_ purging, voltammograms of pyocyanin and methyl viologen displayed diminished oxidation (forward) peaks and altered reduction (backwards) peaks in air (Fig. 5d-f). This matches previous literature which describes the ability of pyocyanin and methyl viologen to generate reactive oxygen species(*56*). Current shifts were also observed in voltammograms recorded in LB in the absence of redox inducers, with a distinctive reduction peak being observed at −439 mV vs SHE (Supplementary Fig. 8), suggesting this is the peak of oxygen reduction.

With these results we demonstrate the activation of the SoxR-P*sox*S system by various classes of redox inducers across a wide range of E_m_ values. Phenazines like pyocyanin and quinones like DHNA have an additional advantage in that they can be naturally synthesised. *E. coli* naturally synthesises DHNA(*57*) and has also been genetic engineered to produce pyocyanin using genes from the *phz* operon of *Pseudomonas*(*58*). Other electrogenic bacteria are also known to produce a variety of other electron mediators which could also act as redox inducers(*25*).

### Electrochemical Control of Gene Expression under Aerobic Conditions

Previous electrogenetic systems have utilised bespoke bioelectrochemical cells and devices in order to perform electrochemical control of gene expression(*22, 26*–*29*). Whilst functional, these *ad hoc* approaches lack standardisation, preventing their use in different electrogenetic systems as well as hindering reproducibility between research groups. We have therefore assembled a bioelectrochemical device specifically designed for use in electrogenetic systems.

This device consists of a stack of acrylic blocks and polydimethylsiloxane (PDMS) gaskets with cylindrical cavities for fastening with stainless-steel screws. Electrically conductive materials serve as working and counter electrode materials with solid acrylic blocks placed at each end of the device. A Nafion® membrane placed at the middle of the stack delimits the working and counter chambers, with magnetic stir bars providing mixing in each chamber (Fig. 6a; Supplementary Fig. 9). The modular nature of the device allows for it to be reconfigured for use with different electrode materials or chamber volumes, meaning the device can be optimised for different applications. The system can be operated in 2- or 3- or 4-electrode mode by introducing one or more reference electrodes. In this study a 3-electrode set-up was used with carbon paper working and counter electrodes and a Ag/AgCl reference electrode, with each chamber having a volume of 30 mL.

**Fig 6:**
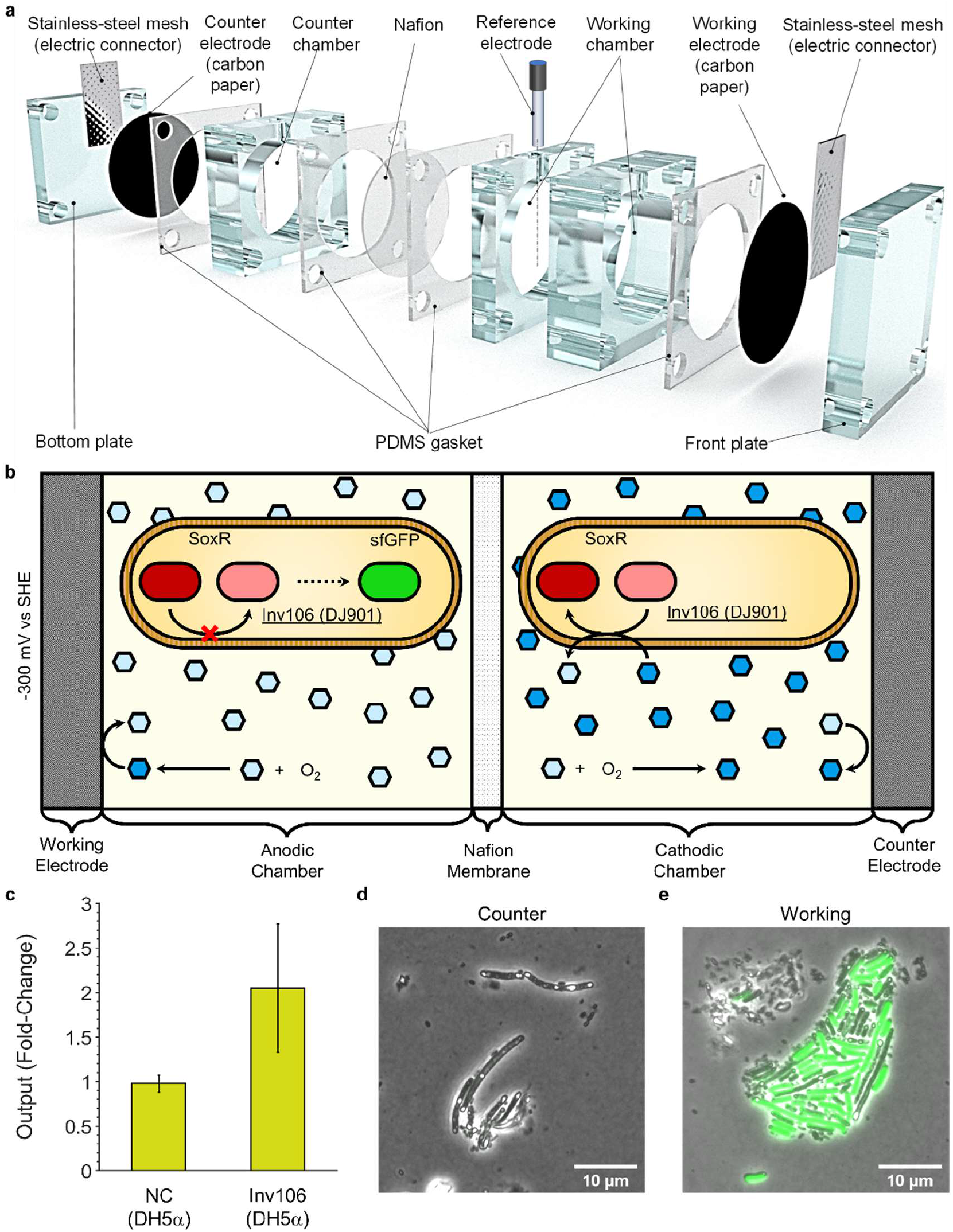
Electrochemical Activation of Gene Expression in Aerobic Conditions. **a**, Diagram of the modular bioelectrochemical device developed for performing electrochemical induction of gene expression. **b**, Schematic of device operation for electrochemical activation of gene expression. The working electrode was held at a potential of −300 mV vs SHE in order to reduce pyocyanin, preventing oxidation of SoxR in Inv106 (DH5α) cells and thereby activating expression of sfGFP in the anodic chamber. **c**, Gene expression change in cells between the working and counter chambers measured as a fold-change. Datapoints represent the mean from three biological replicates with error bars depicting standard deviation (n = 3). **d-e**, Confocal fluorescence micrographs of Inv106 (DH5α) grown in the cathodic and anodic chambers, then immobilised on agarose pads during imaging. sfGFP fluorescence (λ_excitation_ = 490 nm) in green is overlaid on brightfield images. NC, negative control.

With this device we performed electrochemical activation of gene expression in aerobic conditions, utilising pyocyanin as a redox inducer due to its maximal activation of the SoxR-P*sox*S system (Fig. 5b), lack of cytotoxicity with Inv106 (DH5α) (Supplementary Fig. 6e), and its previously recorded long-term stability in bioelectrochemical systems(*30*). Inv106 (DH5α) cells (Fig. 4d) were grown in both chambers of the device in LB electrolyte supplemented with 10 µM pyocyanin under a pyocyanin-reducing applied bias potential of −300 mV vs SHE at the working electrode. If the rate of pyocyanin reduction by the working electrode exceeds the rate of pyocyanin oxidation by oxygen, sfGFP expression is activated in cells within the working chamber by maintaining SoxR in a more reduced state. Alternatively, in the counter chamber pyocyanin should be maintained in an oxidised state by the electrode and oxygen, leading to SoxR activation and suppression of sfGFP expression (Fig. 6b).

Under these conditions Inv106 (DH5α) exhibited a 2.27-fold increase in sfGFP expression in the working chamber relative to the counter chamber, whereas the corresponding negative control (Supplementary Table 3) construct exhibited no significant difference in sfGFP expression between the two chambers (Fig. 6c; Supplementary Fig. 10). This fold-change closely matches those reported for a different *E. coli* electrogenetic system in aerobic conditions(*29*), despite the SoxR-P*sox*S here allowing for a much simpler electrogenetic system without the use of specialised gold-binding strains, a co-culture or minimal media.

The fold-change observed for electrochemical induction of Inv106 (DH5α) was much smaller than that recorded in dose response measurements (Fig. 5f), suggesting that complete reduction of pyocyanin in the anodic chamber was not achieved. Confocal fluorescence microscopy of Inv106 (DH5α) cells from the anodic and cathodic chamber also revealed that whilst cells in the anodic chamber clearly exhibited much higher sfGFP fluorescence, some cells appeared to have reduced fluorescence, suggesting certain heterogeneity within the population (Fig. 6d-e; Supplementary Fig. 11). Regardless, these results demonstrate the first example of electrochemical activation of gene expression in reducing conditions, as well as a proof-of-concept for electrochemical control of gene expression using the SoxR-P*sox*S system in aerobic conditions.

## Discussion

Chemical and optogenetic inducible gene expression systems have been widely applied in molecular biology research as well as being used in various biotechnological devices. Whilst providing effective spatiotemporal control(*22, 26*) and improved integration in bioelectronic devices(*4*), use of electrochemical inducible gene expression systems has been limited due to their reduced performance and dependence on anaerobic conditions or co-cultures. Our results demonstrate significant improvements to the performance of the SoxR-P*sox*S electrochemically inducible gene expression system, providing biological responses comparable to commonly used chemically inducible systems such as the AraC and LasR systems(*46*). A library of redox-responsive promoters allows for rational design of electrogenetic circuits(*31*), and a proof-of-concept for electrochemical gene activation in aerobic conditions is also presented. These promoters can be used to improve the previously developed eCRISPR system for multiplexed control of expression of multiple genes (*25*), as well as facilitating the construction of complex multi-layered logic devices(*59*).

Future research should focus on the improvement of the electrochemical set-up used, as this appears to be the limiting factor in device performance. Optimal performance will be achieved with a device that can provide bulk oxidation or reduction of a stable electron mediator. The modular bioelectrochemical device presented allows for electrodes of different structures and materials to be tested, such as the various biocompatible carbon-based electrodes suitable that have previously been developed for microbial fuel cells(*60*). Identification of suitable redox inducers that exhibit cell permeability, long term stability, low cytotoxicity, minimal oxygen reactivity and the ability to activate the SoxR-P*sox*S system strongly would provide further improvement of the device. Electrochemical limitations could also be overcome through the use of quorum sensing systems and positive feedback circuits to propagate gene expression throughout a culture(*22, 26*). Bacteria could also be engineered to express enzymes that biosynthesise redox inducers; *E. coli* and *Pseudomonas* species have previously been engineered to produce non-endogenous phenazines^65,^(*61*). Genomic integration of electrogenetic constructs will also prevent the requirement for antibiotics which are prone to electrochemical degradation(*62*).

The electrogenetic devices presented here can be easily adapted and integrated for use in a variety of bioelectronics, including medical and environmental biosensors(*4*). The demonstration of electrochemical control of gene expression in relatively large volume cultures (Fig. 6c) also suggests an application in large bioreactors, where low light penetration prevents the use of optogenetic systems. The SoxR-P*sox*S system is also conserved across a diverse range of bacteria(*63*), with electrochemically inducible gene expression with the system being demonstrated in both *E. coli* and *Salmonella enterica*(*26*). Alternative redox-sensing transcription factors to SoxR have been identified in plants(*64*) and animals(*65*). The standard electrogenetic device design we have described here (Fig. 1a) can be applied to these different organisms. Furthermore, the use of the BASIC assembly standard in this study facilitates easy automation of electrogenetic circuit assembly(*38*), aiding the optimisation of new electrogenetic systems.

The lack of available tools for constructing electrogenetic systems has severely limited their development. The optimised toolset we have developed here promises to expediate and expand the development of future electrogenetic systems for a myriad of applications.

## Methods and Materials

### Chemicals and Reagents

Primers and DNA parts (gBlocks™) were synthesized by Integrated DNA Technologies. PCR reactions were carried out using Phusion High-Fidelity DNA Polymerase from Thermo Fisher. Blunt end ligations were performed using the Thermo Fisher CloneJET PCR Cloning Kit. Enzyme digestions were performed using high-fidelity restriction enzymes purchased from New England Biolabs (NEB). DNA ligations were performed using Promega T4 ligase. Redox inducers and other chemicals and materials were purchased from Merck, unless otherwise stated. BASIC linkers were obtained from Biolegio and magnetic DNA purification was done using AMPure XP magnetic DNA purification kits. All organic redox inducers were stored as stock solutions at −20 °C: methyl viologen dichloride hydrate (100 mM in LB), pyocyanin (50 mM in DMSO), DHNA (1 M in DMSO), Riboflavin (1 mM in LB), hydrogen peroxide (1 M in LB).

### Bacterial Strains, Plasmids and Media

The *E. coli* strains used were DH5α (fhuA2 lacΔU169 phoA glnV44 Φ80’ lacZ ΔM15 gyrA96 recA1 relA1 endA1 thi-1 hsdR17), purchased from NEB, and DJ901 (Δ(argF-lac)169 λ-IN(rrnD-rrnE)1 rpsL179(strR) zjc-2205::Tn10kan Δ(soxS-soxR)566), purchased from the Coli Genetic Stock Center (CGSC). *E. coli* strains were cultivated in LB medium and LB-agar plates at 37 °C with corresponding antibiotics at the following concentrations: kanamycin (50 μg/ml), ampicillin (50 μg/ml), chloramphenicol (25 μg/ml). All plasmids contain pMB1 origin of replication with either kanamycin or chloramphenicol resistance, and mScarlet dropout-cassette at the insertion site. For plate reader experiments shown in Supplementary Figure 8, cells were grown in M9 medium (1 × M9 salts, 0.4 % glucose, 2 mM MgSO_4_, 100 µM CaCl_2_) supplemented with 0.2 % casamino acids and 100 mM MOPS, as described in (*22*).

### BASIC DNA Assembly

All part plasmids were assembled by performing blunt end ligations of gBlocks™ into pJET1.2/blunt vectors. All construct plasmids were assembled using BASIC (Biopart Assembly Standard for Idempotent Cloning) assembly(*66*). Parts are listed in Supplementary Table 3, Constructs in Supplementary Table 3 and BASIC linkers in Supplementary Table 4. Plasmid maps are available on GitHub (https://github.com/JLawrence96/ElectrogeneticsToolset/tree/DNA)

### Mutant Library Generation

The promoter library was constructed using PCR mutagenesis with Phusion High-Fidelity DNA Polymerase. Divergent 5’ phosphorylated primers carrying the mutations were used to amplify the whole plasmid and introducer the mutations using the standard Phusion DNA polymerase protocol from NEB. Mutagenesis primers are listed in Supplementary Table 5). Reverse mutagenesis primers were used to mutate the −35 site while forward mutagenesis primers were used to mutate the −10 site. The DNA product was purified using PCR clean-up with QIAquick® PCR Purification Kit, followed by DpnI digestion (NEB) to cut methylated template DNA. Plasmids were re-ligated using Promega T4 before heat-shock transformation into *E. coli* DH5α.

### Chemical Heat-shock E.coli Transformation

Competent cell stocks of *E. coli* DH5α and DJ901 were prepared by the Inoue method(*67*) and stored at −80 °C. Heat-shock transformation was performed by defrosting 50 µl of competent DH5α cells, and immediately adding this to a PCR tube containing 5 µl of plasmid DNA. This mixture was incubated on ice for 20 min before transferring to a thermocycler for heat-shock. Heat-shock was performed with a protocol consisting of 20 min a 4 °C, 45 s at 42 °C and 2 min at 4 °C. Following heat-shock, 200 µL of SOC broth pre-warmed to 37 °C was added. Recovery of transformed culture was performed at 37 °C for 1 hr before 100 µL was plated on LB-agar supplemented with the appropriate antibiotic.

### Stationary Phase Measurements

Glycerol stock of strains containing the constructs and control plasmids were streaked on LB-agar antibiotic plates and incubated overnight at 37 °C. Plates were stored at 4°C. Single colonies were picked from these plates and inoculated into 5 mL LB supplemented with the appropriate antibiotic and incubated overnight at 37 °C. 2 µL of overnight cultures were diluted in 198 µL of LB + antibiotics with or without pyocyanin to an OD_600_ of approximately 0.1. LB media containing pyocyanin was created by diluting pyocyanin stock solutions in LB so that diluted overnight cultures had the desired working concentration of pyocyanin (either 2.5 or 10 µM). These cultures were transferred to wells of a 96-well microplate (Costar) which was sealed with Breathe-Easy® sealing membrane which was placed in a Synergy HT microplate reader (BioTek) and incubated at 37 °C with orbital shaking at 1000 rpm for 14 h. After incubation, endpoint sfGFP fluorescence (excitation, 485 nm; emission, 528 nm; gain 40) and OD_600_ measurements were taken with a Synergy HT microplate reader (BioTek). For measuring the upstream activity, RFP measurements (excitation, 590 nm; emission, 645 nm; gain 70) were also taken. All measurements were taken from the bottom. Outputs were expressed in units of RPU. This was calculated by subtracting blank fluorescence and OD_600_ measurements from the data (which were recorded from measurements of the media condition without cells), before dividing fluorescence measurements by OD_600_ measurements and expressing them as a ratio relative to the same measurements recorded from a strain harbouring an RPU standard plasmid in LB medium. This plasmid contained the same fluorophore being expressed from a J23101 promoter with the same standardised design used for the other promoters in this study (*38, 41*) (Supplementary Table 3).

For plate reader experiments shown in Supplementary Figure 8, cells were treated identically except M9 medium (Bacterial Strains, Plasmids and Media) was used instead of LB medium.

### Response Function Measurements

Microplates were prepared identically to how they were prepared for stationary phase measurements, apart from overnight cultures were diluted in LB or M9 media + antibiotics containing different redox inducers to a range of different redox inducer concentrations. Response curves were calculated by a previously developed method(*14*) in which experimental data was fitted to a straight line for non-responsive constructs, to equation [1] for Uni-P*sox*S and activator devices, and to equation [2] for inverter devices:

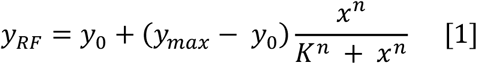

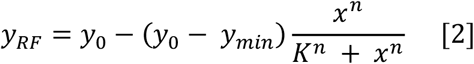

where *y*_*RF*_ is the fitted response function. The fixed parameters were: *x*, the concentration of redox inducer; *y*_0_, the RPU with no redox inducer added; *y*_*max*_, the largest achieved RPU value across all concentrations of redox inducer. The fitted parameters were: *K*, the sensitivity; *n*, the cooperativity.

Fitting was performed in MATLAB using a custom script utilising the fminsearch function. Response functions were fit by the least-squares method to minimise the Sum of Squared Estimate of Errors (SSE) between the data and the model, as detailed in equation [3]:

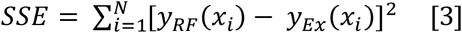

where *y*_*RF*_(*x*_*i*_) is the response function RPU value and *y*_*Ex*_(*x*_*i*_) is the experimental RPU value for a given concentration of redox inducer *x*_*i*_, with *N* being the total number of redox inducer concentrations tested.

The dynamic rage was calculated from experimental data, as detailed in equation [4] for Uni-P*sox*S and activator devices, and to equation [5] for inverter devices:

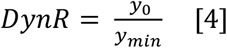

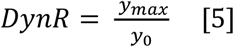

The R^2^ was also calculated for each model to determine the quality of the fit, as detailed in equation [6]:

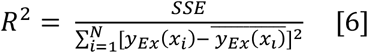

All experimental and calculated values of response functions are listed in Supplementary Table 1. All Code is available on Github (https://github.com/JLawrence96/ElectrogeneticsToolset/tree/Code)

### Cyclic Voltammetry

Cyclic voltammetry was performed with a PalmSens EmStat3 Blue. A 20 mL glass vial was used as an electrochemical cell with a glassy carbon working electrode, platinum mesh counter electrode and a Ag/AgCl reference electrode (Supplementary Fig. 6a). Voltammograms were recorded from 5 mL of LB medium alone, or supplemented with either 1 mM of methyl viologen, 100 µM of pyocyanin or 1 mM DHNA. The cell was heated to 37 °C using a hot plate. Three scans were performed for each sample between using a scan rate of 10 mV s^−1^. Scans were recorded both with and without purging. Purging was performed for 30 mins with N_2_ gas bubbled through a long needle, with said needle being placed in the headspace of the vial when the scan was recorded to maintain anoxic conditions. The third scan for each experiment was recorded, with the corresponding scan of LB medium over the same potential being subtracted from it. E_m_ values were calculated from the purged condition from the potential lying equidistant between the oxidation and reduction peaks. Potentials were converted from mV vs Ag/AgCl to mV vs SHE by addition of 200 mV.

### Bioelectrochemical Device Fabrication

The modular bioelectrochemical device used in this study consisted of a stack of acrylic blocks (Engineering & Design Plastics Ltd, UK), polydimethylsiloxane (PDMS) gaskets, carbon paper electrodes (Fuel Cell Store, Texas, US), Nafion® 117 membrane and stainless-steel mesh electrical connectors (MeshDirect Ltd, UK).

From front to back, the stack was formed of: solid acrylic blocks (Supplementary Fig. 9a) serving as a front plate followed by a carbon paper disk working electrode (Supplementary Fig. 9e) with a stainless-steel mesh used as an electrical connector (Supplementary Fig. 9g) and a PDMS gasket (Supplementary Fig. 9b). Then, two acrylic blocks with cylindrical central cavity (Supplementary Fig. 9c-d) were placed next to each other forming the working chamber. A reference electrode was placed in the working chamber by a vertical port present in one of the acrylic blocks. The working chamber was followed by a disk of Nafion® 117 membrane (Supplementary Fig. 9f) sandwiched between two PDMS gasket (Supplementary Fig. 9f). The device continued with an acrylic block with a cylindrical central cavity (Supplementary Fig. 9b) forming the cathodic chamber followed by a PDMS gasket (Supplementary Fig. 9f), a carbon paper disk counter electrode (Supplementary Fig. 9e) and a stainless-steel mesh used as an electrical connector (Supplementary Fig. 9g). The device was then completed with a solid acrylic block for the back plate (Supplementary Fig. 9a). All those components were fastened together with fours stainless-steel screws. The complete device is shown in Fig. 6a and Supplementary Fig. 9h. Magnetic stir bars providing mixing were placed in each chamber and the device was placed atop a multi-plate stirrer (Svelp Scientific, Italy), with rubber stoppers being used to plug any cavities during experiments.

### Electrochemical Induction of Gene Expression

Overnight cultures were diluted in LB + chloramphenicol to a final OD_600_ of 0.05. Pyocyanin stock was added to cultures to achieve a final concentration of 10 µM. 30 mL of culture was loaded into each chamber of the bioelectrochemical device which was then sealed with rubber plugs, placed on a magnetic stirrer in an incubator set to 37 °C and connected to a PalmSens EmStat3 Blue. Chronoamperometry was performed for 16 hr with an applied bias potential of −500 mV vs Ag/AgCl (−300 mV vs SHE) and a sampling rate of 1 s^−1^. Following this cultures from each chamber were visualised by confocal fluorescence microscopy and then concentrated 4-fold before measuring OD_600_ using a X UV-visible spectrophotometer and fluorescence using an Edinburgh Instruments FS5 Spectrofluorometer (excitation, 485 nm; excitation bandwidth, 1 nm; emission, 528 nm; emission bandwidth, 0.3 nm). Output was measured by dividing fluorescence measurements by OD_600_ measurements obtained using a Varian Cary 50 Bio UV-vis spectrometer.

### Confocal Fluorescence Microscopy

Live Inv106 (DH5α) cells that were grown in the working and counter chambers of the bioelectrochemical device were immobilised on agarose pads made of 1% low-melt agarose solution in water(*68*). Images were acquired using a Nikon Eclipse Ti widefield microscope with a Nikon objective lens (Plan APO, 100×/1.45 oil) and a Hamamatsu C11440, ORCA Flash 4.0 camera. Brightfield and fluorescence of sfGFP (λ_excitation_ = 490 nm) images were taken. Images were processed using NIS Elements Viewer and Image J software.

## Acknowledgements

We thank Prof. Richard Kitney, Prof. Tom Ellis, Dr. Robert Bradley and Dr. Ari Dwijayanti for helpful discussions.

## Author contributions

J.M.L, Y.Y & R.L-A designed the study. J.M.L & Y.Y performed the experiments. Pixcell iGEM Team performed preliminary studies. P.B. constructed the bioelectrochemical device. A.S. assisted with plate reader experiments. M.S assisted with BASIC assembly. L.T.W. performed the fluorescence microscopy. J.M.L developed the code, analysed the data and wrote the manuscript with input from all authors.

## Data and materials availability

All data needed to evaluate the conclusions in the paper are present in the paper and/or the Supplementary Materials. All input data, code and plasmid maps from this study are available on GitHub here: https://github.com/JLawrence96/ElectrogeneticsToolset.

