## Supplementary Material for "A modular toolset for electrogenetics"

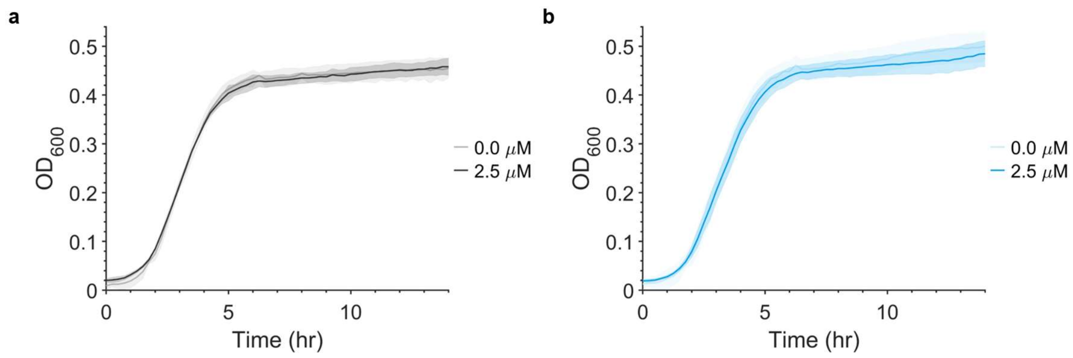


**Supplementary Fig. 1:** Cytotoxicity of promoter constructs.
**a,** Negative control (DR) and **b,** Uni-P*sox*S construct (DR) growth curves with and without 2.5 µM pyocyanin. Solid lines represent the mean from three biological replicates with shaded areas depicting standard deviation of the mean (n = 3).

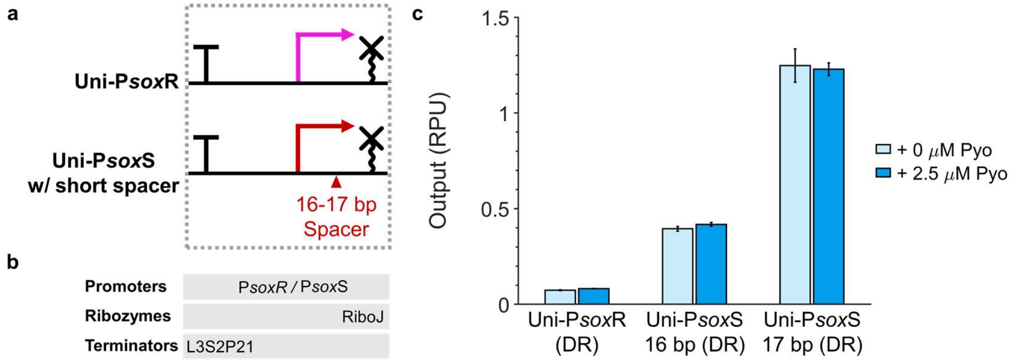


**Supplementary Fig. 2:** Screening the activity of alternative P*sox*R and P*sox*S variants.
**a,** Architectures of engineered Uni-P*sox*R promoter and Uni-P*sox*S promoters with shortened spacers. Promoters were tested using the genetic circuit in Fig. 2c. **b,** List of genetic parts used. **c,** Downstream activity of engineered promoters with and without 2.5 μM pyocyanin. Datapoints represent the mean from three biological replicates with error bars depicting standard deviation of the mean (n = 3). DR, downstream reporter; RPU, relative promoter units.

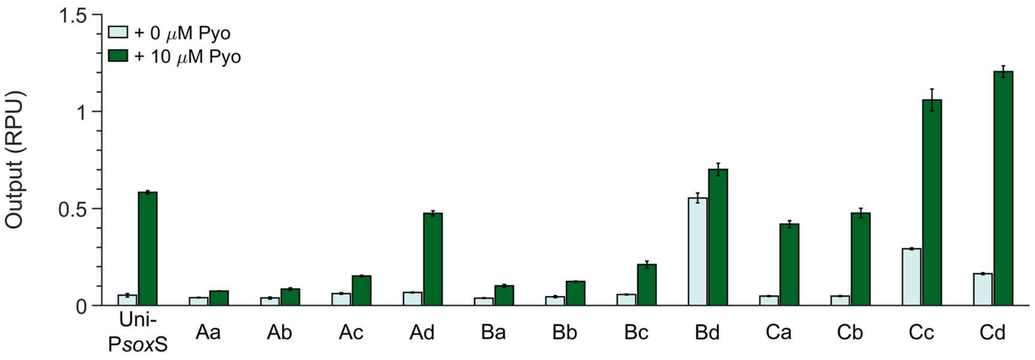


**Supplementary Fig. 3:** Screening Uni-P*sox*S promoter library variants.
Downstream activity of engineered promoters with and without 2.5 μM pyocyanin. Promoters were tested using the genetic circuit in Fig. 2c. Upper case characters refer to the -35 sites and lowercase characters the -10 sites listed in Fig. 3a. Datapoints represent the mean from three biological replicates with error bars depicting standard deviation of the mean (n = 3). RPU, relative promoter units.

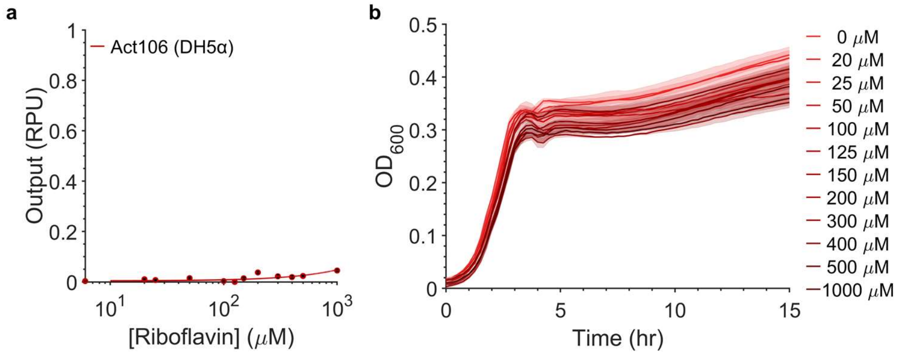


**Supplementary Fig. 4:** Electrogenetic circuit performance with riboflavin
**a,** Response function of Act106 (DJ901) with riboflavin. Datapoints represent the mean from three biological replicates with error bars depicting standard deviation of the mean (n = 3). **b,** Growth curve of Act106 (DJ901) in increasing concentrations of riboflavin. Solid lines represent the mean from three biological replicates with shaded areas depicting standard deviation of the mean (n = 3).

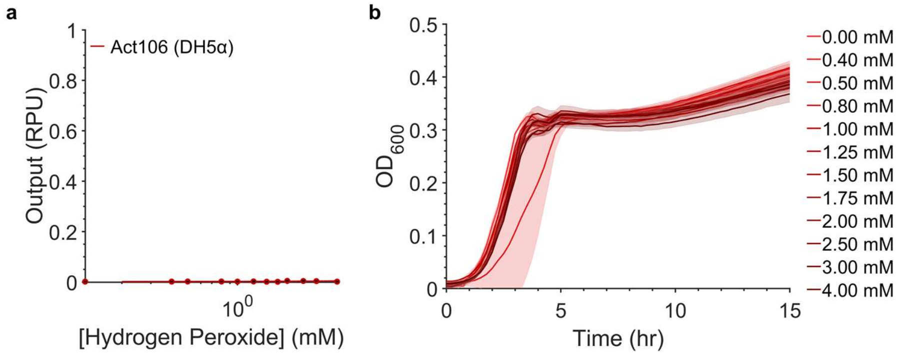


**Supplementary Fig. 5:** Electrogenetic circuit performance with hydrogen peroxide
**a,** Response function of Act106 (DJ901) with hydrogen peroxide. Datapoints represent the mean from three biological replicates with error bars depicting standard deviation of the mean (n = 3). **b,** Growth curve of Act106 (DJ901) in increasing concentrations of hydrogen peroxide. Solid lines represent the mean from three biological replicates with shaded areas depicting standard deviation of the mean (n = 3).

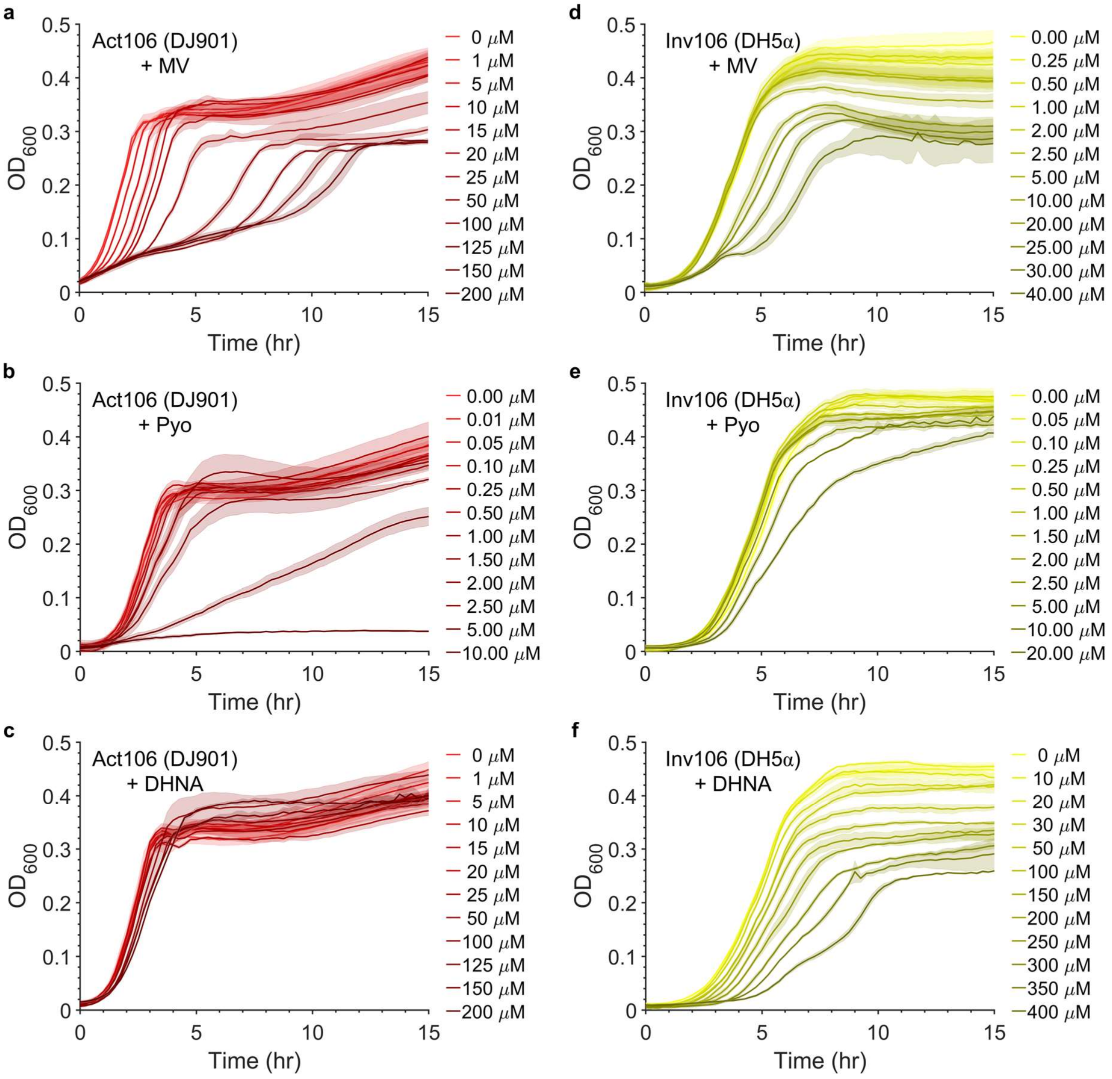


**Supplementary Fig. 6:** Cytotoxicity of electrogenetic circuits with diverse redox inducers.
**a-c,** Act106 (DJ901) and **d-f,** Inv106 (DH5α) growth curves in increasing concentrations of methyl viologen, pyocyanin and DHNA. Solid lines represent the mean from three biological replicates with shaded areas depicting standard deviation of the mean (n = 3).

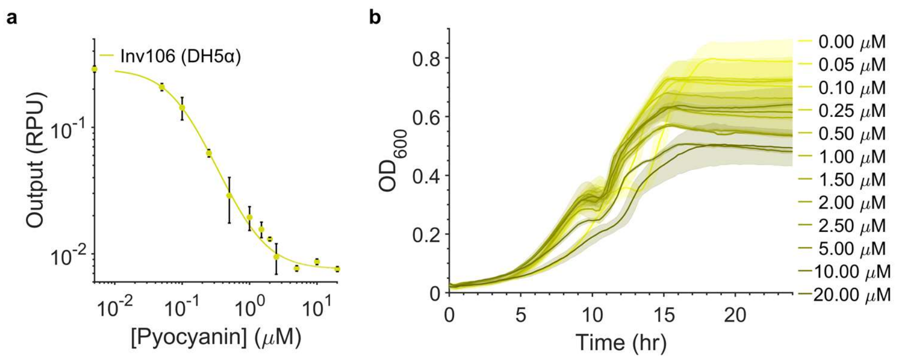


**Supplementary Fig. 7:** Electrogenetic circuit performance in minimal media
**a,** Response function of Inv106 (DH5α) with pyocyanin in M9 media. Datapoints represent the mean from three biological replicates with error bars depicting standard deviation of the mean (n = 3). **b,** Growth curve of Inv106 (DH5α) in increasing concentrations of pyocyanin in M9 media. Solid lines represent the mean from three biological replicates with shaded areas depicting standard deviation of the mean (n = 3).

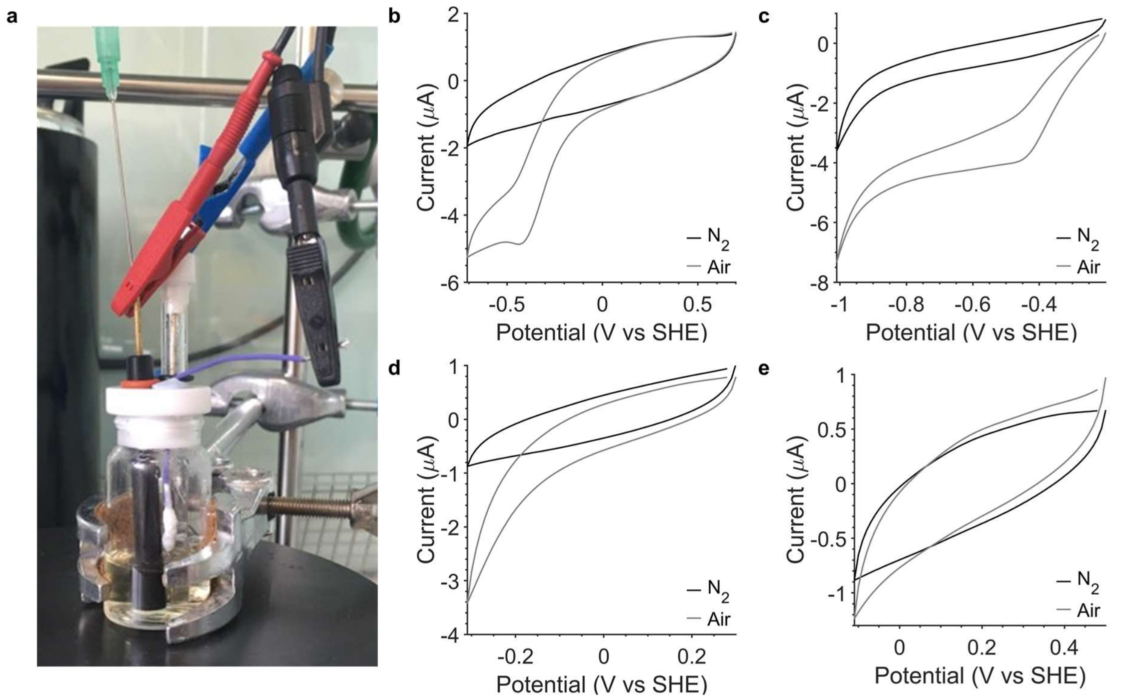


**Supplementary Fig. 8:** Cyclic voltammetry of LB media
**a,** Photograph of the bioelectrochemical cell used for cyclic voltammetry experiments. Red connects to the glassy carbon working electrode, black to the platinum mesh counter electrode and blue to the Ag/AgCl reference electrode. **b,** Cyclic voltammograms of LB medium over a large potential range. **c-e,** Cyclic voltammograms performed over the same potential ranges as Fig. 5b-d. The third scan of each voltammogram was recorded using a scan rate of 10 mV $s^{-1}$. Voltammograms were recorded both in air and in electrolyte purged with $N_2$ gas for 30 mins (to remove oxygen) before measurements were recorded with an $N_2$ gas stream being maintained in the headspace.

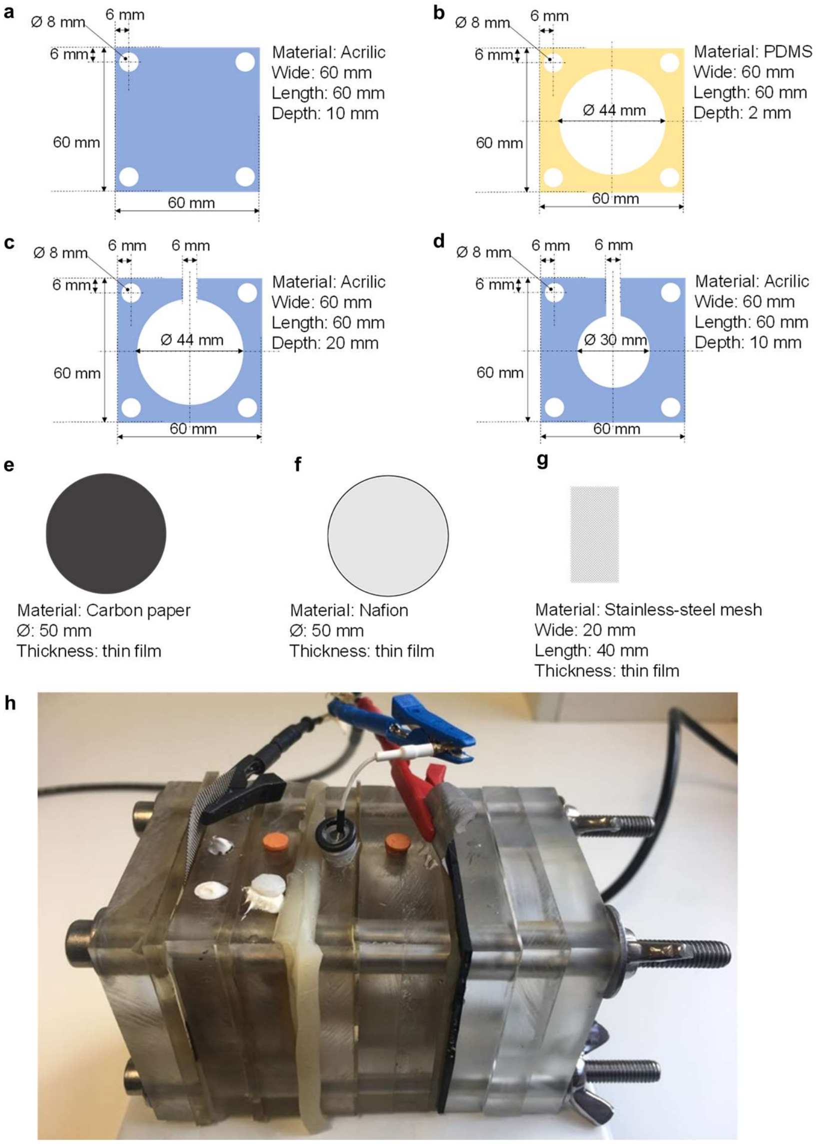


**Supplementary Fig. 9:** Bioelectrochemical device design
Dimensions and materials of **a,** front/back plates, **b,** gaskets, **c,** working and counter electrode chambers, **d,** reference chamber, **e,** working and counter electrodes, **f,** Nafion membrane and **g,** electric connectors. **h,** Photograph of the assembled device. Red connects to the working electrode, black to the counter electrode and blue to the reference electrode.

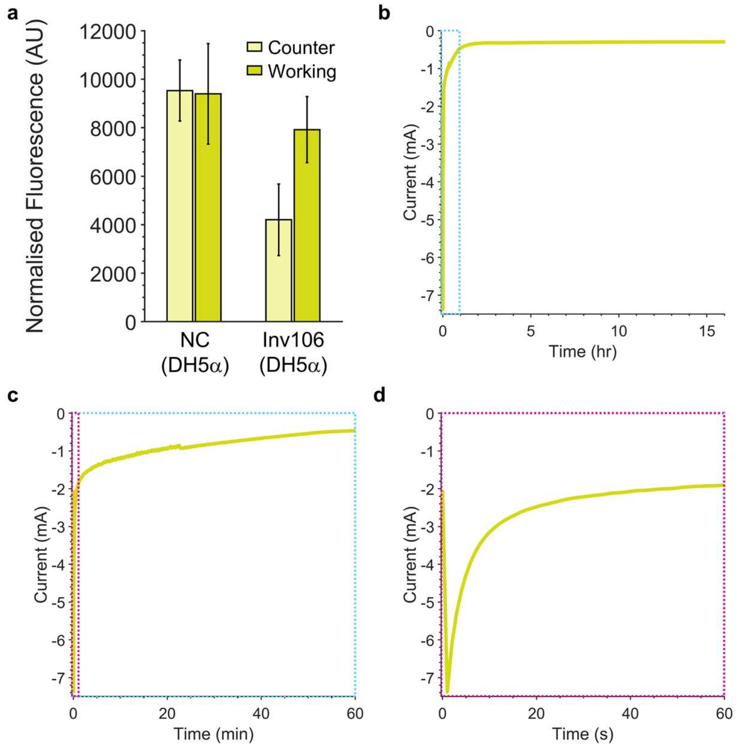


**Supplementary Fig. 10:** Chronoamperometry of Inv106 (DH5α) in the bioelectrochemical device
**a,** Gene expression change in cells between the working and counter chambers measured by cell-normalised fluorescence. Datapoints represent the mean from three biological replicates with error bars depicting standard deviation of the mean (n = 3). **b,** Representative chronoamperometric scan recorded during electrochemical activation of gene expression of an Inv106 (DH5α) culture in LB medium supplemented with chloramphenicol and 10 μM pyocyanin. Scans were performed over 16 hours with an applied bias potential of -300 mV vs SHE at the working electrode and a sampling rate of 1 $s^{-1}$. **c-d,** Zoom in of the first hour and first minute of the scan.

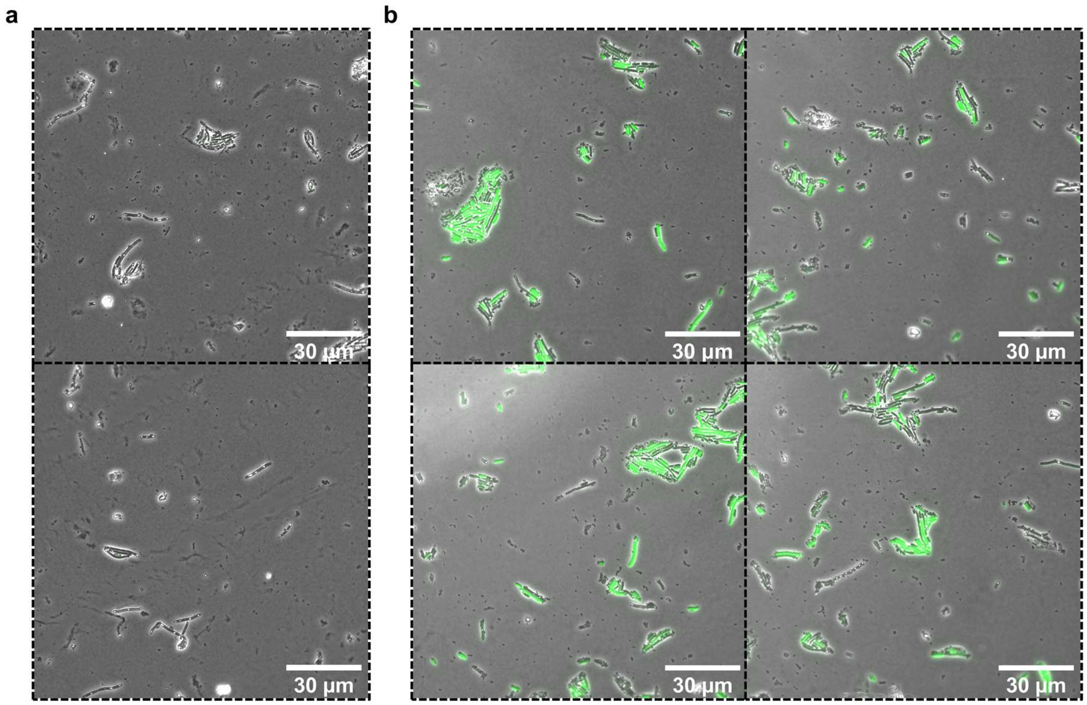


**Supplementary Fig. 11:** Imaging electrochemical activation of gene expression
Uncropped confocal fluorescence micrographs of Inv106 (DH5α) grown in the **a,** counter electrode and **b,** working electrode chamber. sfGFP fluorescence in green is overlaid on brightfield images. All micrographs were recorded from a single biological replicate.

| Construct | Inducer | Max Concentration (μM) | $y_0$ (RPU) | $y_{min}/y_{max}$ (RPU) | $DynR$ | $n$ | $K$ (μM) | $R^2$ |
|---|---|---|---|---|---|---|---|---|
| Uni-P*sox*S | Pyocyanin | 40 | 0.079 | 0.884 | 11.19 | 1.22 | 4.29 | 0.979 |
| Act106 (DJ901) | Methyl Viologen | 200 | 0.105 | 2.492 | 23.68 | 1.04 | 22.12 | 0.892 |
| Inv106 (DH5α) | Methyl Viologen | 40 | 0.839 | 0.053 | 15.70 | 1.70 | 4.76 | 0.996 |
| Act106 (DJ901) | Pyocyanin | 10 | 0.105 | 4.824 | 46.06 | 1.88 | 1.46 | 0.998 |
| Inv106 (DH5α) | Pyocyanin | 20 | 0.996 | 0.020 | 49.97 | 1.37 | 0.57 | 0.998 |
| Act106 (DJ901) | DHNA | 200 | 0.099 | 2.116 | 21.39 | 2.57 | 64.38 | 0.977 |
| Inv106 (DH5α) | DHNA | 400 | 0.977 | 0.248 | 3.93 | 2.55 | 138.82 | 0.991 |
| Inv106 (DH5α) | Pyocyanin (M9) | 20 | 0.289 | 0.008 | 38.15 | 1.44 | 0.09 | 1.000 |

**Supplementary Table 1:** Response function parameters

| Plasmid Index | Name | Function | Antibiotic | Part Length (bp) |
|---|---|---|---|---|
| pPEG.000 | pBR322-BASIC Kan$^R$ | Construct Acceptor | Kanamycin | 3698 |
| pPEG.001 | pBR322-BASIC Cm$^R$ | Construct Acceptor | Chloramphenicol | 3554 |
| pPEG.100 | Native P*sox*R/S | Promoter | Ampicillin | 63 |
| pPEG.101 | Bi-P*sox*S | Promoter | Ampicillin | 219 |
| pPEG.102 | Uni-P*sox*S | Promoter | Ampicillin | 209 |
| pPEG.103 | Uni-P*sox*S-Aa | Promoter | Ampicillin | 209 |
| pPEG.104 | Uni-P*sox*S-Ab | Promoter | Ampicillin | 209 |
| pPEG.105 | Uni-P*sox*S-Ac | Promoter | Ampicillin | 209 |
| pPEG.106 | Uni-P*sox*S-Ad | Promoter | Ampicillin | 209 |
| pPEG.107 | Uni-P*sox*S-Ba | Promoter | Ampicillin | 209 |
| pPEG.108 | Uni-P*sox*S-Bb | Promoter | Ampicillin | 209 |
| pPEG.109 | Uni-P*sox*S-Bc | Promoter | Ampicillin | 209 |
| pPEG.110 | Uni-P*sox*S-Bd | Promoter | Ampicillin | 209 |
| pPEG.111 | Uni-P*sox*S-Ca | Promoter | Ampicillin | 209 |
| pPEG.112 | Uni-P*sox*S-Cb | Promoter | Ampicillin | 209 |
| pPEG.113 | Uni-P*sox*S-Cc | Promoter | Ampicillin | 209 |
| pPEG.114 | Uni-P*sox*S-Cd | Promoter | Ampicillin | 209 |
| pPEG.115 | Uni-P*sox*R | Promoter | Ampicillin | 208 |
| pPEG.116 | Uni-P*sox*S 16 bp | Promoter | Ampicillin | 206 |
| pPEG.117 | Uni-P*sox*S 17 bp | Promoter | Ampicillin | 207 |
| pPEG.118 | J23101 | Promoter | Ampicillin | 191 |
| pPEG.119 | J23101_Inverted | Promoter | Ampicillin | 191 |
| pPEG.120 | J23105 | Promoter | Ampicillin | 187 |
| pPEG.121 | J23106 | Promoter | Ampicillin | 191 |
| pPEG.122 | P*phl*F | Promoter | Ampicillin | 238 |
| pPEG.200 | GFP | CDS | Ampicillin | 714 |
| pPEG.201 | RFP_Inverted | RBS+CDS+Terminator | Ampicillin | 925 |
| pPEG.202 | SoxR | CDS | Ampicillin | 465 |
| pPEG.203 | PhlF | CDS | Ampicillin | 603 |
| pPEG.300 | B14 | Terminator | Ampicillin | 95 |

**Supplementary Table 2:** BASIC parts list
Full plasmid maps are contained within the supplementary files.

| Plasmid Index | Name | Backbone | Antibiotic | Parts & Linkers |
|---|---|---|---|---|
| pCEG.000 | NC (Promoter & Act) | pPEG.000 | Kanamycin | LMP - pPEG.300 - LMS |
| pCEG.001 | Native (DR + UR) | pPEG.000 | Kanamycin | LMP - pPEG.201 - L4 - pPEG.100 - U2RBS3 - pPEG.200 - LMS |
| pCEG.002 | Native (DR) | pPEG.000 | Kanamycin | LMP - pPEG.100 - U2RBS3 - pPEG.200 - LMS |
| pCEG.003 | Bi-P*sox*S (DR + UR) | pPEG.000 | Kanamycin | LMP - pPEG.201 - L4 - pPEG.101 - U2RBS3 - pPEG.200 - LMS |
| pCEG.004 | Bi-P*sox* (DR) | pPEG.000 | Kanamycin | LMP - pPEG.101 - U2RBS3 - pPEG.200 - LMS |
| pCEG.005 | Uni-P*sox*S (DR + UR) | pPEG.000 | Kanamycin | LMP - pPEG.201 - L4 - pPEG.102 - U2RBS3 - pPEG.200 - LMS |
| pCEG.006 | Uni-P*sox*S (DR) | pPEG.000 | Kanamycin | LMP - pPEG.102 - U2RBS3 - pPEG.200 - LMS |
| pCEG.007 | Uni-P*sox*R (DR) | pPEG.000 | Kanamycin | LMP - pPEG.115 - U2RBS3 - pPEG.200 - LMS |
| pCEG.008 | Uni-P*sox*S 16 bp (DR) | pPEG.000 | Kanamycin | LMP - pPEG.116 - U2RBS3 - pPEG.200 - LMS |
| pCEG.009 | Uni-P*sox*S 17 bp (DR) | pPEG.000 | Kanamycin | LMP - pPEG.117 - U2RBS3 - pPEG.200 - LMS |
| pCEG.010 | Act105 | pPEG.000 | Kanamycin | LMP - pPEG.120 - U3RBS2 - pPEG.202 - L4 - pPEG.102 - U2RBS3 - pPEG.200 - LMS |
| pCEG.011 | Act106 | pPEG.000 | Kanamycin | LMP - pPEG.121 - U3RBS2 - pPEG.202 - L4 - pPEG.102 - U2RBS3 - pPEG.200 - LMS |
| pCEG.012 | NC (Inv) | pPEG.001 | Chloramphenicol | LMP - pPEG.122 - U1RBS3 - pPEG.200 - L1 - pPEG.300 - L2 - pPEG.118 - U3RBS1 -pPEG.203 - LMS |
| pCEG.013 | Inv105 | pPEG.001 | Chloramphenicol | LMP - pPEG.122 - U1RBS3 - pPEG.200 - L1 - pPEG.300 - L2 - pPEG.102 - U3RBS1 -pPEG.203 - L4 -pPEG.120 - U2RBS2 -pPEG.202 - LMS |
| pCEG.014 | Inv106 | pPEG.001 | Chloramphenicol | LMP - pPEG.122 - U1RBS3 - pPEG.200 - L1 - pPEG.300 - L2 - pPEG.102 - U3RBS1 -pPEG.203 - L4 -pPEG.121 - U2RBS2 -pPEG.202 - |
| pCEG.015 | RPU_Calibrant (GFP) | pPEG.000 | Kanamycin | LMP - pPEG.118 - U2RBS3 - pPEG.200 - LMS |
| pCEG.016 | RPU_Calibrant (RFP) | pPEG.000 | Kanamycin | LMP - pPEG.201 - L1 - pPEG.119 - LMS |

**Supplementary Table 3:** BASIC construct list
Parts are in red, standard BASIC linkers in pink, methylated prefix/suffix linkers in blue and RBS linkers in orange. Full plasmid maps are contained within the supplementary files.

| Name | Linker Type | Sequence (5’-3’) |
|---|---|---|
| L1 | Neutral | CTCGttacttacgacaCTCCGAGACAGTCAGAGGGTAtttattgaactaGTCC |
| L2 | Neutral | CTCGatcggtgtgaaaAGTCAGTATCCAGTCGTGTAGttcttattacctGTCC |
| L4 | Neutral | CTCGagaagtagtgccACAGACAGTATTGCTTACGAGttgatttatcctGTCC |
| LMP | Methylated | CTCGggtaagaactcgCACTTCGTGGAAACACTATTATCtggtgggtctctGTCC |
| LMS | Methylated | CTCGggagacctatcgGTAATAACAGTCCAATCTGGTGTaacttcggaatcGTCC |
| U1RBS3 | RBS | CTCGttgaacaccgtcTCAGGTAAGTATCAGTTGTAAAAagaggagaaataGTCC |
| U2RBS2 | RBS | CTCGtgttactattggCTGAGATAAGGGTAGCAGAAAAAagaggggaaataGTCC |
| U2RBS3 | RBS | CTCGtgttactattggCTGAGATAAGGGTAGCAGAAAAAagaggagaaataGTCC |
| U3RBS1 | RBS | CTCGgtatctcgtggtCTGACGGTAAAATCTATTGTAATcacacaggactaGTCC |
| U3RBS2 | RBS | CTCGgtatctcgtggtCTGACGGTAAAATCTATTGTAAAagaggggaaataGTCC |

**Supplementary Table 4:** BASIC linker list
Sequences are annealed linker sequences after BASIC assembly. Scar regions are in red and upper case. Overhang regions are in black and upper case and adapter regions are in pink and lower case, unless they have other functionality. Prefix and suffix regions of methylated linkers are in blue and RBSs in orange.

| Name | Orientation | Function | Sequence (5'-3') |
|---|---|---|---|
| SP1_pJET_F | Forward | Sequencing | CGACTCACTATAGGGAGAGCGGC |
| SP2_pJET_R | Reverse | Sequencing | AAGAACATCGATTTTCCATGGCAG |
| SP3_pBR322_F | Forward | Sequencing | GGCGGCGGATTTGTCCTAC |
| SP4_pBR322_R | Reverse | Sequencing | GGACCCCTGGATTCTCACC |
| SP5_RFP_F | Forward | Sequencing | GGCTCGGGAGACCTATCGATCTT |
| SP6_RFP_R | Reverse | Sequencing | GCCATCTAGTATTTCTCCTCTTT |
| CP1_Psox_F | Forward | Cloning | TCTGGTGGGTCTCTGTCCAAATCGCTTTACCTCAAGTTAACTT |
| CP2_ Psox_R | Reverse | Cloning | CGATAGGTCTCCCGAGCCCAGTTCGTTAATTCATCTGTTGG |
| CP3_SoxR_F | Forward | Cloning | TCTGGTGGGTCTCTGTCCTTAGTTTTGTTCATCTTCCAGCA |
| CP4_SoxR_R | Reverse | Cloning | CGATAGGTCTCCCGAGCCATGGAAAAGAAATTACCCCG |
| MP1_35A | Reverse | Mutagenesis | Pi-TAACTTGAGGTAAAGCGATTTAGATTTTCCAGAGTAGGCAC |
| MP2_35B | Reverse | Mutagenesis | Pi-TAACTTGAGGGCAAGCGATTTAGATTTTCCAGAGTAGGCAC |
| MP3_35C | Reverse | Mutagenesis | Pi-TAACTTGAGGTCAAGCGATTTAGATTTTCCAGAGTAGGCAC |
| MP4_10a | Forward | Mutagenesis | Pi-ACTTGAGGAATTATGTTCCCCAACAGATGAATAGCTG |
| MP5_10b | Forward | Mutagenesis | Pi-ACTTGAGGAATTATAAACCCCAACAGATGAATAGCTG |
| MP6_10c | Forward | Mutagenesis | Pi-ACTTGAGGAATAATAATCCCCAACAGATGAATAGCTG |
| MP7_10d | Forward | Mutagenesis | Pi-ACTTGAGGAATTATAATCCCCAACAGATGAATAGCTG |

**Supplementary Table 5:** Primer list
Overhang regions are in blue with "Pi" denoting the primers are 5' phosphorylated.
